# Cortico-thalamic excitatory-inhibitory dynamics generates *α*-wave transient patterns encoding anesthetic brain states

**DOI:** 10.64898/2025.12.25.696413

**Authors:** François David, Christophe Sun, Pierre-Olivier Michel, Jérémie Sibille, Vincenzo Crunelli, Nathalie Rouach, David Holcman

## Abstract

The neuronal circuits generating *α* frequency (8-13Hz) oscillatory patterns during anesthesia are poorly understood, making their use difficult in predictive medicine. Here, we combine large-scale in vivo recordings of thalamocortical neuronal ensembles in anesthetized mice with a minimal neural mass model. We report non-rhythmic, weakly synchronized firing of thalamic and cortical neurons correlated with *α* oscillatory patterns, indicating an emergent network rather than cellular dynamics. The associated imbalance between the firing of excitatory and inhibitory cortical neurons, together with an increase in firing fluctuations, favors larger amplitudes *α* oscillations, whereas changes in thalamic neuron output control *α* oscillation frequency. A neural mass model, comprising reciprocally connected excitatory and inhibitory cortical and thalamic neurons well reproduces the *α* oscillation patterns observed in vivo at increasing anesthetic doses. These results provide a direct link between EEG *α* oscillations during anesthesia and their underlying network/cellular dynamics which are well captured by a computational model that is clinically relevant for anesthetized brain state monitoring.

**Highlights:**

- *α* transient patterns during anesthesia emerge from thalamocortical and cortical excitatory–inhibitory (E–I) network dynamics.
- *α* pattern frequency and amplitude reflect depth of anesthesia, dynamic thalamocortical equilibrium and E–I imbalance.
- Emergence of spindle-like patterns at multiple frequency defines newly characterized brain-state during anesthesia
- A mass-model reproduces the transient nature and spectral diversity of anesthetic-induced *α* oscillations, linking neural circuit mechanisms to EEG output.

## Introduction

Discontinuous oscillations in the *α* frequency band (8-13 Hz) are commonly observed in the EEG during clinical anesthesia with different agents, including propofol or halogenic gasses [1]. These transient events are characterized by an increasing amplitude (waxing) at their start and a decreasing amplitude (waning) at their end, which has led to their designation as *α*-spindles, though here we will retain the more generic term of *α* oscillation for a single motif instead of spindle.

It is known that thalamic neurons can entrain layer 4 cortical neurons at *α* frequency during relaxed wakefulness [2, 3] while the cortex can also sustain *α* oscillatory activity when isolated from the thalamus [4]. Although these thalamic and cortical *α* oscillations have a waveform and frequency similar to those observed during anesthesia [5, 6], no detailed study has investigated the mechanisms underlying these transient *α* oscillatory patterns during anesthesia. Moreover, the bouts of *α* activity observed during iso-electric suppression in very deep anesthesia [7] have not been fully characterized [8]. This information is critical not only for our mechanistic understanding, but also for improving clinical anesthesia monitoring and real time prediction by computational models [9, 10]. In particular, models of *α* oscillations have often been restricted to a cellular view or to a stable oscillation [2, 11, 12, 13, 14, 15], which do not reflect the waxing and waning or the diversity of alpha oscillations. Alternatively, some mean-field models, which abstract the biophysical complexity and neuronal spike mechanisms, better accounted for the emergence principles and fragmentation of oscillations in the *α* frequency band [16, 17] but lack a direct comparison to the physiological and neuronal correlates of anesthesia.

Here, we investigated the firing dynamics of large ensembles of single thalamocortical (TC) neurons together with cortical excitatory (i.e. regular spiking, RS) and inhibitory (i.e. fast spiking, FS) neurons in mice under different depths of anesthesia. These data were used to construct a neuronal mass-model driven by noise [18, 19] that could define the characteristic elements of anesthetic-induced *α* oscillations. We found that non-rhythmic and weakly synchronized firing of thalamic and cortical neurons underlie *α* oscillatory patterns. The associated imbalance of excitatory and inhibitory cortical neurons firing and an increase in their firing fluctuations modulate *α* oscillations amplitude, whereas changes in thalamic neuron output control *α* oscillations frequency. Our neuronal mass model, comprising reciprocally connected excitatory and inhibitory cortical and thalamic neurons well reproduces the different characteristics of *α* oscillation patterns observed in vivo at increasing anesthetic doses, and provides a mechanistic framework for integrating dose-dependent modulation of *α* oscillations and their relationship to evolving brain states.

## Results

### Spike-train statistics of thalamic and cortical neurons during *α* oscillations

To investigate how cellular firing dynamics contributes to the neuronal population activity during anesthetic-induced oscillations in the *α* frequency band, we recorded the local field potential (LFP) simultaneously with spikes from individual neurons using 384-channel Neuropixels probes located horizontally in layer 4 of the occipital cortex of mice anesthetized with isoflurane (Fig. 1A-B)(see Methods). In particular, we isolated the firing of excitatory thalamocortical (TC) axons (as a proxy of TC neuron activity), of cortical excitatory regular spiking (RS) neurons and cortical inhibitory fast spiking (FS) neurons (Fig. 1B)(see Methods). The power spectrum of the LFP (Fig. 1C) shows a convex shape, suggesting a deviation from a power law EEG [20, 21]. Notably, although *α* activity in anesthetized humans is centered around 10 Hz, *α* oscillation frequency in mice can span between 4 Hz [22, 23] during light anesthesia and up to 16 Hz when these EEG waveforms emerge from the burst-suppression associated with deep anesthesia[24].

**Figure 1.**
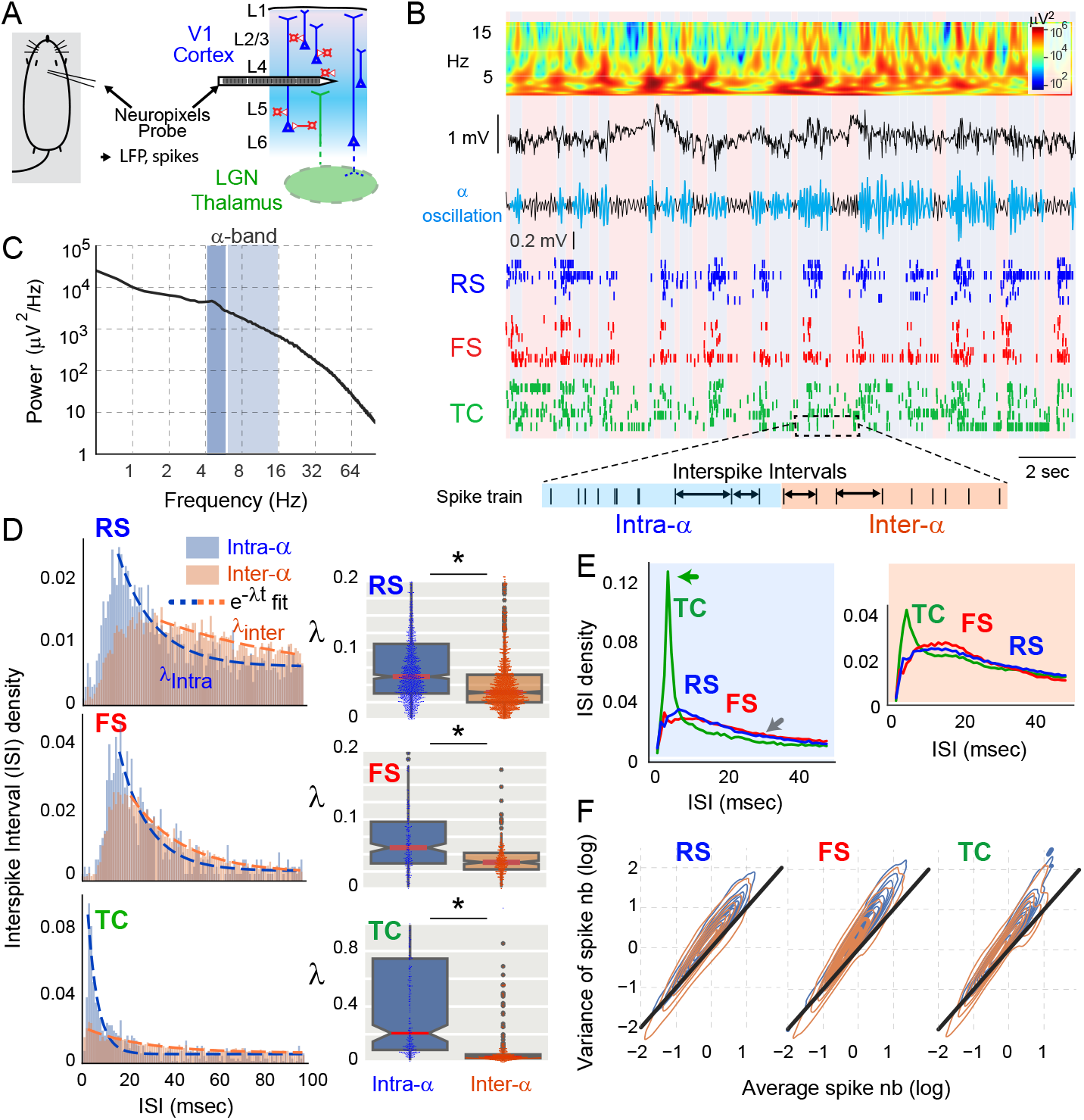
Stochastic neuronal firing during *α* oscillations. **(A)** Schematic drawing of recordings in anesthetized mice. The 384-channel Neuropixels probe positioned in layer 4 of the primary visual cortex collects the local field potential (LFP) and spikes of local cortical neurons and thalamocortical (TC) axons. **(B)** From top to bottom: LFP scalogram (colormap), raw LFP (black), band-pass (6-16 Hz) filtered LFP overlaid by segmented *α* oscillations (blue) and spike time rasters of 20 excitatory cortical neurons (RS), 20 inhibitory cortical neurons (FS) and 20 excitatory thalamocortical (TC) neurons all overlaid over Intra-*α* (vertical blue bars) and Inter-*α*(vertical orange bars) events. **(C)** LFP power spectrum. In dark and light blue, the low (4-6Hz) and high (6-16Hz) *α* frequency bands, respectively. **(D)** Left: Interspike Interval (ISI) density distributions for a typical RS, FS and TC neuron for Intra- *α* (blue) and Inter-*α* (orange) oscillations. Dashed lines represent the fit with an exponential function *a* exp(−*λt*). Right: *λ*-values (red: median; edges: quartiles; crosses: outliers) of each analyzed neuron (dots) for Intra-*α* and Inter-*α* oscillations (RS, n=1227, FS, n=228, TC, n=290 neurons). ∗ : *p <*0.05. **(E)** Average ISI distribution density of all RS, FS and TC neurons for Intra-*α* (blue background) and Inter-*α* (red background) oscillations. Note 1) that the sharp peak (green arrow) at 2 ms in the TC distribution corresponds to high-frequency bursts of action potentials, 2) the overlap of the RS and FS distribution (grey arrow) **(F)** Density distribution (density lines) of variance versus mean number of spikes in 100 sequences taken from Intra- (blue) and Inter- (orange) *α* oscillations for RS (slope=1.21 vs 1.15 for Intra-*α* and Inter-*α* oscillations, respectively), FS (slope=1.30 vs 1.29) and TC (slope=1.24 vs 1.30). The two distributions of the three neuron types are distinct from the diagonal black line (equivalent to a Poisson process with slope=1) and between each other (*p <* 0.05).

We next analyzed the variability of the spike trains to determine whether their statistics were random, i.e. characterized by a Poisson process. We found that the distribution of the interspike intervals (ISI) measured by fitting the tail of the distribution with an exponential rate (*λ*) differed between intra- and inter- *α* oscillatory events (Intra-*α* and Inter-*α* respectively) (RS: *λ*_*Intra*_ = 0.061*s*^−1^, *λ*_*Inter*_ = 0.039*s*^−1^, n=1227, *p <* 0.01, Wilcoxon test), (FS: *λ*_*Intra*_ = 0.059*s*^−1^, *λ*_*Inter*_ = 0.038*s*^−1^, n=228, *p <* 0.01),(TC: *λ*_*Intra*_ = 0.21*s*^−1^, *λ*_*Inter*_ = 0.035*s*^−1^, n=290, *p <* 0.01, Wilcoxon test)(n=4 mice)(Fig. 1 D). We also observed that the distribution tails of the RS, FS and TC neurons overlapped both for Intra- and Inter-*α* oscillatory events (Fig. 1E). However, for short ISIs the TC neuron distribution differed from that of RS and FS neurons since in the former population there is a large number of high-frequency bursts of action potentials during anesthesia and *α* oscillations (Fig. 1E) [25, 26].

Finally, to estimate the possible deviation of these statistics from a Poisson process, we measured the number of spikes and represented their mean vs. variance over 100 samples [27] in Intra- and Inter-*α* periods. In both cases, the density distribution of the mean vs. the variance differed from a pure Poisson process (diagonal: mean=variance)(p≪ 0.01,*T >* 18 for all cases, t-tests run against a slope of one) (Fig. 1F). Moreover, though significant (*p <* 0.01, Ancova tests, F=300 for TC, 1480 for RS, 86 for FS), the difference between Intra- and Inter- *α*-events is very small (RS slope=1.21 vs 1.15 Intra-*α* and Inter-*α* oscillations respectively; FS slope=1.30 vs 1.29; and TC slope=1.24 vs 1.30), suggesting that spike train statistics are closely related for Intra-*α* and Inter-*α* oscillations.

In summary, the spike distributions for both intra- and inter-*α* oscillatory events approach a random Poisson distribution well with distinct rates. However, these rates are similar between FS and RS neurons, but differ for TC neurons.

### Emergence of excitatory and inhibitory population synchrony during the waxing and waning of *α* oscillations

The relationship between the firing of individual neurons and the waxing and waning dynamics of *α* oscillations remains unclear. We observed a clear increase and decrease in the firing at the start (waxing) and end (waning), respectively, of the *α* epochs coupled with a respective increase and decrease in the amplitude of the LFP (Fig. 2A). To quantify these firing changes, we first identified the LFP minima of the *α* oscillation waveform to be used as a time reference point (Fig. 2B, top) so that the spike histogram could be compared when spikes were aligned with the first, second, and third cycles from the beginning of an *α* oscillation event (when its amplitude is increasing) or with last three cycles of the end of the oscillation (when its amplitude is decreasing), as shown in Fig. 2B.

**Figure 2.**
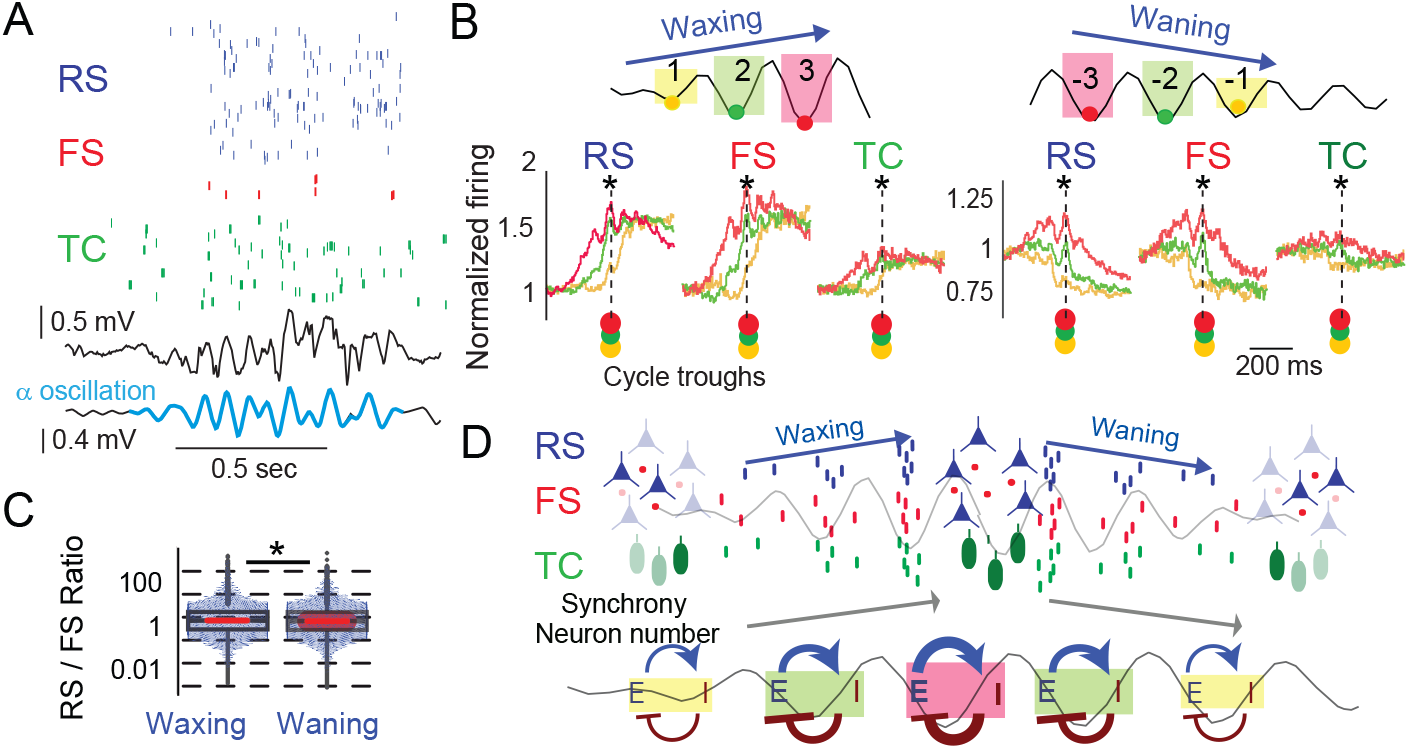
Synchronous excitatory and inhibitory neuronal assemblies emerge during *α* oscillations. **(A)** From top to bottom: raster plot of RS, FS and TC neurons, LFP and filtered LFP during an *α*-oscillation (blue) (RS, n = 28, FS, n = 6, TC, n = 39 neurons). **(B)** Comparison of neuronal synchrony in the first 3 and the last 3 *α* oscillation cycles for the three types of neurons (RS, n = 1227, FS, n = 228, TC, n = 290). The firing histograms centered at the first and last three *α* oscillation troughs indicated synchrony increase and decrease at the beginning (waxing) and end (waning) of the *α* oscillation, respectively (∗: *p <* 0.05 between the 1st vs. 2nd and the 2nd vs. 3rd LFP troughs (color dots)). **(C)** Firing ratio of RS and FS neurons during waxing and waning (∗ *p <*0.05). **(D)** Suggested schematic population dynamics of excitatory and inhibitory networks during waxing and waning. As neurons are more synchronous and fire more while the *α* oscillation grows (first row), the coupling strength increases (second row) (arrows indicate excitatory and inhibitory connections).

We found that for all three neuronal populations, the troughs associated with the waxing and waning of the *α* oscillation are associated with a ramping increase and decrease, respectively, of population firing (*p* ≪ 0.05, w *>* 5000, Wilcoxon test, RS, n = 1227, FS, n = 228, TC, n = 290). Notably, the peak of the population firing was well aligned with the troughs of the LFP (Fig. 2B). Moreover, we found only a small (2.5%) difference (median=0.74 (95% CI:0.077,7.4) vs 0.72 (95% CI:0.072,7.7), *p <* 0.01, Wilcoxon test n=6190 *α* oscillations) in the firing ratios of RS and FS neurons between the waxing and the waning phases (Fig. 2C) indicating that the relative firing of the excitatory and inhibitory cortical populations remains almost unaltered during these phases. On the basis of these results, we suggest that the *α* oscillation pattern could result from mutual interactions between inhibitory and excitatory neurons, with more neurons being entrained and synchronized when the *α* oscillation amplitude increases (Fig. 2D).

### Stochastic neural-mass models generate *α* oscillations with typical waxing and waning patterns

Since the increased coupling between the excitatory and inhibitory network is driven by similarly random noise (Fig. 1D-F), we used these characteristics to build a neural mass model (Fig. 3A) based on recurrent interactions between an excitatory and an inhibitory population. The model consists of an excitatory population (E) that excites an inhibitory population (I) with strength *ω*_1_ and inhibits itself with a self-inhibition decay rate *λ*, and an inhibitory population (I), that inhibits the excitatory network with strength *ω*_2_ and itself with a self-inhibition decay rate *λ* (Fig. 3A) (method eq.2). In this model we tested whether random activity driven by a Brownian noise (*σ*) was sufficient to generate *α* oscillations and whether changes in the neuronal population activities as observed experimentally could account for the characteristic waxing and waning patterns of these oscillations.

**Figure 3.**
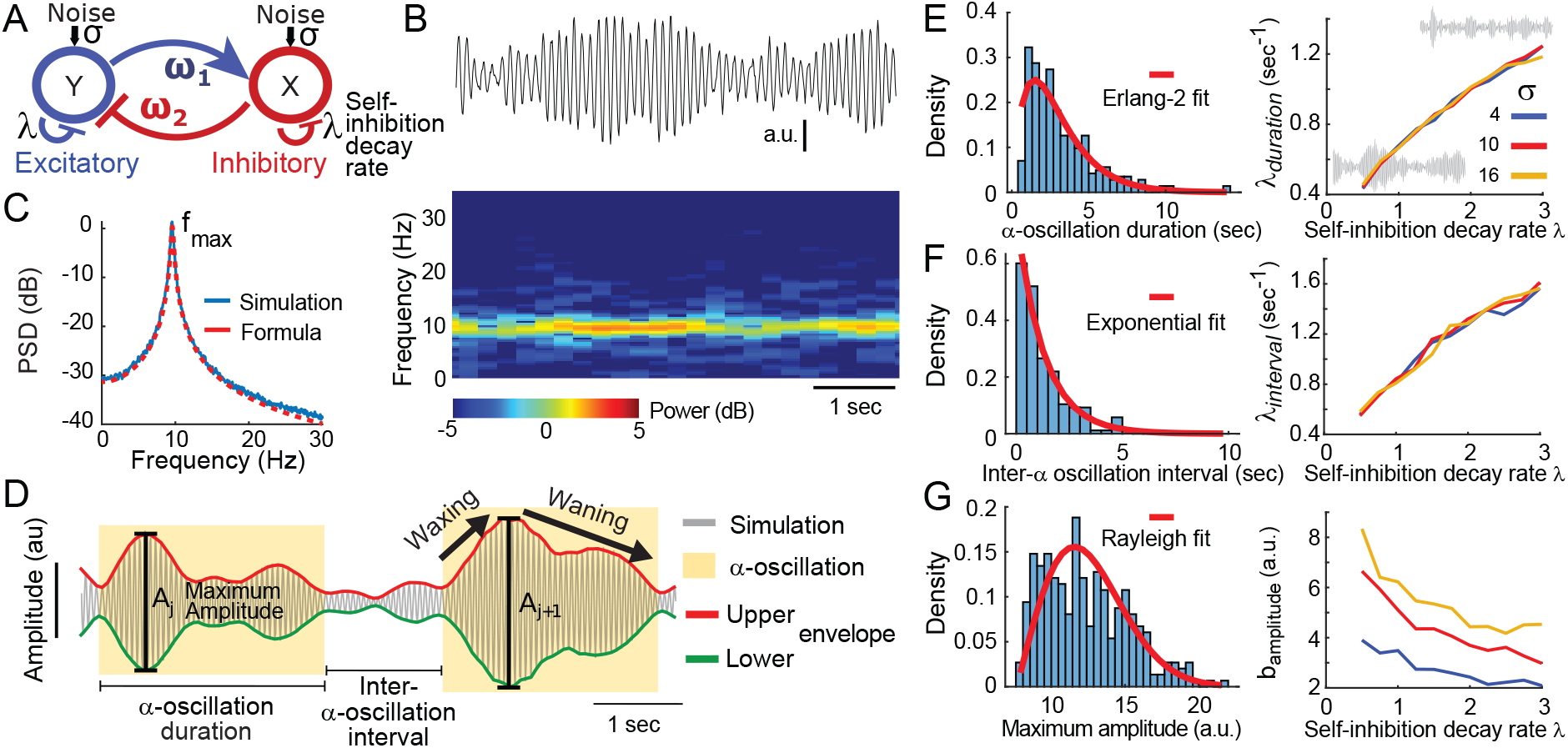
*α* oscillation events emerge from the stochastic process model with two interacting mean field populations. **(A)** Synaptic interactions are modeled through a mean-field excitatory (E) and inhibitory (I) neuronal network with direct (*ω*_1_, *ω*_2_) and feedback self-inhibition (*λ*) interactions driven by noise *σ* forming a two-dimensional Ornstein-Uhlenbeck (OU) process. **(B)** The first component *x*(*t*) of a model realization (amplitude in arbitrary unit (a.u.)) with parameters *λ* = 1, *ω*_1_ = *ω*_2_ = 60, *σ* = 8 in eq. 2 and below its spectrogram (amplitude color coded). **(C)** The power spectrum density estimated from simulations (blue) and computed analytically (red) obtained from eq.(7). **(D)** *α* oscillation event segmentation from one realization of the model and definition of its features. The upper (red) and lower (green) lines interpolated between local maxima and minima define the envelope. **(E)** Left: distribution of *α* oscillation event durations over a 20-min simulation and its fit (red) using an Erlang-2 law. Right: Evolution of the fit parameter *λ*_*duration*_ with respect to the self-inhibition decay rate *λ* for different OU noise amplitudes (*σ* = 4, 10, 16). Inset grey traces: 10-sec simulation example with *λ* = 0.5 (bottom left trace) or *λ* = 3 (top right trace) and *ω* = 60, *σ* = 8. **(F)** Left: distribution of Inter-*α* oscillation intervals and its fit (red) using an exponential law. Right: evolution of the exponential decay parameter *λ*_*interval*_ with respect to the self-inhibition decay rate *λ* for different noise amplitude. **(G)** Left: distribution of the *α* oscillation maximum amplitude *A* and its fit (red) using a Rayleigh law. Right: evolution of the Rayleigh scale *b* parameter with respect to the self-inhibition decay rate *λ* for different noise amplitudes.

Numerical simulations of this model revealed the emergence of spontaneous and discrete *α* oscillation events (Fig. 3B) with their power spectrum showing a single peak around a resonance frequency of 10 Hz (Fig. 3C). Next, to characterize the *α* oscillation dynamics generated by the model, we segmented *α* oscillations into single spindle-like events and extracted their individual features i.e. duration, intervals, waxing and waning duration, local maxima and amplitudes of the waveform envelope (Fig. 3D). Based on this segmentation, we fitted the duration distribution with the Erlang-2 function 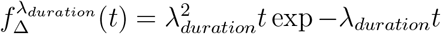 (Fig. 3E left). Interestingly, the oscillation duration distribution (parameterized with the rate *λ*_*duration*_) depends quasi-linearly on the self-inhibition decay rate *λ* (eq. 2), but remains independent of the noise amplitude *σ* (Fig. 3E right). Thus, the higher the self-inhibition decay rate, the shorter the *α* oscillation events. A similar relation to *λ* is observed for the durations of the waxing and waning phases (Fig. S1A-B). Similarly, the distribution of the intervals between *α* oscillation events could be fitted by a single exponential with a decay rate *λ*_*interval*_ proportional to self-inhibition decay rate *λ* (Fig. 3F). Thus, the higher the self-inhibition rate, the shorter the intervals between events, thus favoring a fragmentation of transient *α* oscillations. Lastly, the distribution of maximum amplitude of the filtered signal was fitted with a Rayleigh function 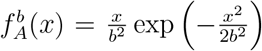 (method eq. 6) where *b* = 4.3 *±* 0.2 (Fig. 3G left). The scale parameter *b* decreases with self-inhibition rate *λ*, while it increases with the amplitude of noise *σ* (Fig. 3G right). In comparison, the filtered signal envelope shape also changes with the inhibition decay rate (Fig. S1C). Thus, the higher the self-inhibition rate, the smaller in amplitude the *α* oscillation patterns and the higher the Brownian noise *σ* the longer the duration of the oscillatory event and the waxing and waning patterns.

In summary, the *α* oscillation events generated by the stochastic model depend on a few parameters (*λ, ω, σ*). Moreover, the individual oscillatory events are widely distributed across the feature dimensions and thus cannot individually be directly used to reflect the state of the E-I network and may carry only little coding information about their generators.

### In vivo *α* oscillation events obey the stochastic model statistic laws

To validate the above model, we applied the same segmentation procedure and statistical analysis to the segments of the LFP recordings corresponding to low (at 0.7% isoflurane) (Fig. 4A-F) and high (1.5% isoflurane) (Fig. 4G-L) dose of anesthetic. For the low doses, the dominant frequency is centered at around 5Hz (Fig. 4B-C) and we obtained a good accuracy for both the duration and the Inter- *α* oscillation intervals 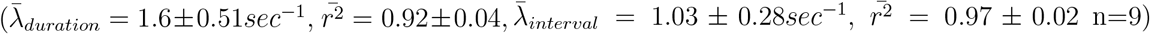. Similarly, an exponential fit explained well the waxing and waning durations 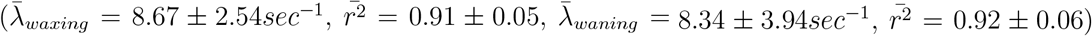 (Fig. 4D, right). In addition, from the segmented event we obtained a mean frequency of 5.48 *±* 0.51 Hz, 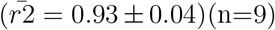(Fig. 4E). Furthermore, the filtered signal maxima could also be well fitted by a Rayleigh distribution (Fig. 4F) as predicted by the statistical laws that characterize the model 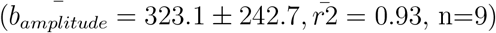.

**Figure 4.**
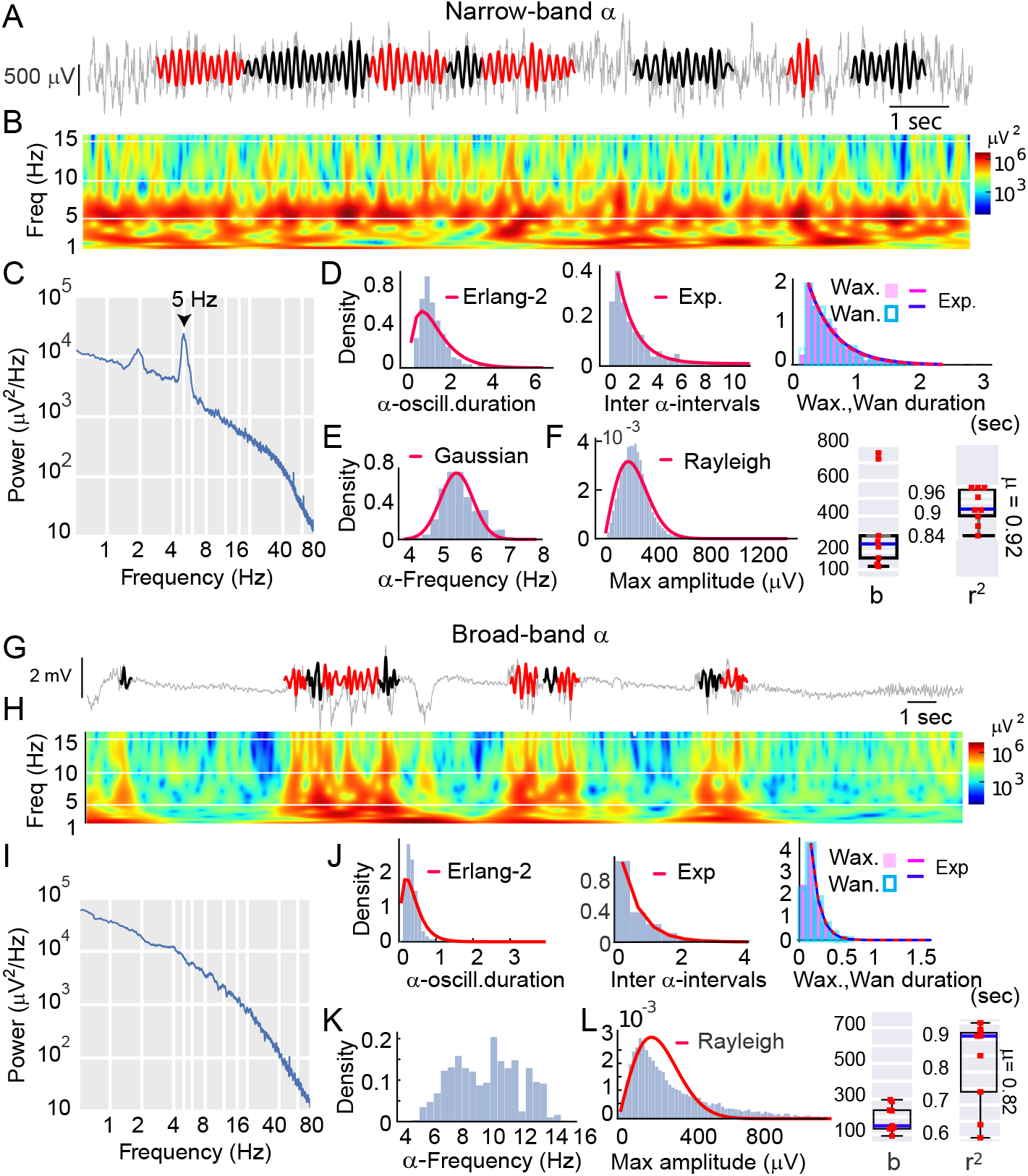
*α* oscillations of different frequency bands emerge at light and deep anesthesia levels. **(A)** Raw LFP with segmented *α* oscillations (overlay in alternating red and black sections) during 0.7% isoflurane anesthesia. **(B)** LFP scalogram showing a dominance of 5 Hz *α* oscillation activity. **(C)** Power spectrum of the LFP over a 1-min period including the LFP trace shown in B. **(D)** Left: Distribution of *α* oscillation durations and its fit (red) using an Erlang-2 law *λ*_*duration*_ = 1.57, *r*^2^ = 0.65. Middle: Distribution of inter *α* oscillation interval durations and its fit (red) using an exponential function *λ*_*interval*_ = 0.65, *r*^2^ = 0.99. Right: distribution of the *α* oscillation waxing (pink) and waning (blue) durations *λ*_*ExpW ax*_ = 3.7, *r*^2^ = 0.99,*λ*_*ExpW an*_ = 65, *r*^2^ = 0.99 respectively. n=1 mouse, n=231 events. **(E)** Distribution of *α* oscillation frequencies and its fit (red) using a normal law *µ* = 5.54, *r*^2^ = 0.94 n=1 mouse. **(F)** Left: distribution of maximum amplitude and its fit (red) using a Rayleigh law *b* = 191.5, *r*^2^ = 0.94 n=1 mouse. Right: Rayleigh scale *b* and fit r^2^ scatter and box plot indicate good fit performance for n=9 mice. **(G-L).** Similar analysis as A-F for a deeper anesthesia level (1.5% isoflurane) *λ*_*duration*_ = 5.0, *r*^2^ = 0.81, *λ*_*interval*_ = 1.62, *r*^2^ = 0.99, *λ*_*W axing*_ = 16.9, *r*^2^ = 0.99,*λ*_*W aning*_ = 15.8, *r*^2^ = 0.99, *b* = 198.9, *r*^2^ = 0.8 n=1 mouse. Population data n=9 mice.

During deeper anesthesia the segmentation revealed different spectrograms enriched towards the 6-16 Hz band while the 4-6 Hz band was decreased (Fig. 4G-I). Similarly to the low anesthetic dose, the duration of the *α* oscillation event, the inter-*α* oscillation intervals, and the waxing and waning were well fitted by the same laws : 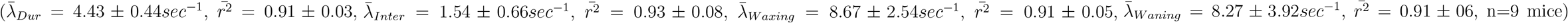. However, the *α* oscillation frequency distribution did not follow a single Gaussian 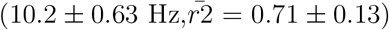 (n=9) (Fig. 4K), and the Rayleigh function was unable to appropriately fit the distribution of local LFP envelope maxima 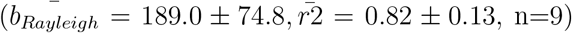, with the *r*^2^ values being significantly smaller than the ones obtained for the narrow band *α* characteristics)(p¡0.01, Mann-Whitney U test, n=9) Fig. 4L). This deviation suggests either a heterogeneous *α* oscillation distribution (see below) or a different *α* oscillation generation mechanism. At this stage, we conclude that the present model statistics well account for the in vivo *α* oscillations at 5Hz.

### Deeper anesthesia boosts *α* oscillation amplitude via a greater Excitation Inhibition imbalance

To investigate how progressive increases in the depth of anesthesia influence the *α* oscillations, we studied changes in LFP amplitude and the underlying E-I balance in cortical networks in mice undergoing stepwise increases in isoflurane dose from 0.5% to 2.1% in steps of 0.2% every 5 minutes (Fig. 5A). Bandpass filtering in the *α*-range (6–16 Hz) revealed a progressive increase in oscillatory amplitude (positive slope = 0.14*µV/s, p <* 0.05, Wilcoxon test, n=9 mice)(Fig. 5B, left, Fig. S2). The *α* oscillation spindle-like events segmentation using our detection procedure (Fig. 5A, boxes) revealed larger-amplitude *α* oscillation at higher isoflurane doses, where the mean event amplitude, calculated in sliding windows of 100 s, increased over time in individual animals (Fig. 5B). This was accompanied by a concomitant significant decrease of the firing rate of the simultaneously recorded RS and FS neurons (Fig. 5C)(*p <* 0.05, Wilcoxon test, n=9). However, extraction of local maxima of RS and FS neuron firing rates indicated an increasing slope for RS neurons and a decreasing slope for FS which are significantly different (Fig. 5C, green fit and right column, *p <* 0.05, Wilcoxon test, n=9). Moreover, the FS neuron firing rate decreased more than that of RS neurons (Fig. 5C, red fit and middle column) as indicated by their significant difference (Fig. 5C, bottom, *p <* 0.05, Wilcoxon test, n=9), suggesting a gradual shift of the E–I balance toward excitation.

**Figure 5.**
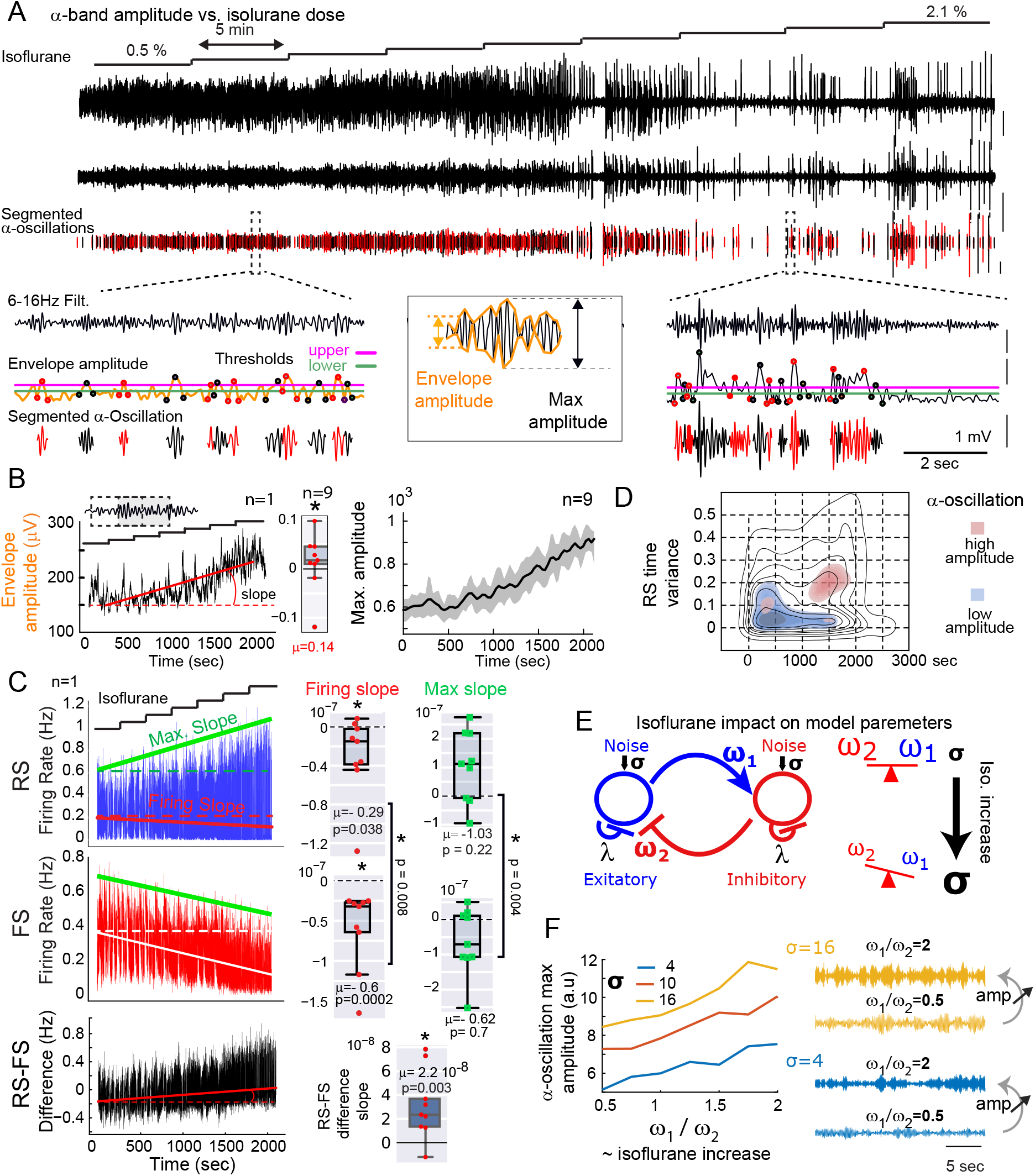
Increasing anesthesia depth enhances *α* oscillation amplitude and shifts the E–I balance towards excitation. **(A)** Experimental protocol showing isoflurane concentration increased from 0.5% to 2.1% in 0.2% steps every 5 minutes. Top to bottom: (1) Raw local field potential (LFP) and (2) band-pass filtered LFP in the *α* band (6–16 Hz) show a progressive increase in oscillation amplitude; (3) Segmented *α* oscillation (waxing–waning events) identified by maxima of the envelope amplitude above an upper threshold (pink) flanked by two minima below a lower threshold (green). Insets illustrate the segmentation procedure at low (left) and high (right) isoflurane doses. **(B)** Left: envelope amplitude (n = 1 mouse) computed over a 100 s sliding window increases over time (red linear fit). Middle: positive slope of the envelope amplitude over time is consistent across 9 mice (Wilcoxon test, *p <* 0.05). Right: *α* oscillation maximal amplitude *±* SEM (gray) (n = 9 mice) increases over the first 2100 s. **(C)** Example (left) and group data (right) for RS and FS firing rate and their difference. Linear fits of the firing rate (red or white) and difference show significant slopes (middle) as isoflurane increases (* *p <* 0.05, n = 9). Linear fits of local maxima (computed on a sliding 100 s window) indicate opposite and significantly different slopes between RS and FS (green) (∗ : *p <* 0.05, n=9). *µ* and p on the group data indicate means and p-values. **(D)** Left: density plots (lines) of RS firing time variance as isoflurane increases over time. The color code indicates *α* oscillations with highest and lowest amplitude quantiles (*q*_0.33_ and *q*_0.66_) (n = 9 mice). **(E)** Model schematic illustrating how increasing isoflurane biases the E–I balance toward excitation and increases noise. **(F)** Model output: *α* oscillation amplitude increases with both the E-I ratio (*ω*_1_*/ω*_2_) and noise amplitude (*σ* color coded).

Because of this effect of isoflurane on the E-I neuronal balance, we next directly correlated *α* oscillation single event amplitude with the firing rate of cortical neurons (Fig. S3). RS neuron firing and the difference between RS and FS firing, but not FS neuron firing were positively correlated with the *α* oscillation event maximum amplitude (Fig. S3). The time variance in the firing time of the RS (but not of the FS neurons) also correlated positively with the *α* oscillation amplitude (Fig. S3). However, the variance in RS firing across RS neurons was not correlated with *α* oscillation amplitude (Fig. S3). Lastly, we found that the temporal variance of RS firing (Fig. S3 top right) also increased, and decreased, with the isoflurane dose for the high, and the low amplitude of the *α* oscillations, respectively (Fig. 5D). This suggests that the increase in amplitude is mainly driven by the proportion of events with a large temporal fluctuation of the RS firing, while lower amplitude events are associated with a small temporal fluctuation of the RS firing. Thus, the increase in *α* oscillation single spindle-like amplitude is driven 1) by the increase in RS and FS population firing differences and 2) by the increase in firing temporal fluctuations in cortical neurons.

To interpret these changes in amplitude, we used our computational model in which the *α* oscillation amplitude depends on the E–I ratio and the level of background noise (Fig. 5E). Increasing the E–I ratio in the model reproduced the experimental increase in *α* oscillation amplitude (Fig. 5F), although the effect of an overall decrease of the level of firing of E and I separately (Fig. S4A-C), that is to decrease the amplitude, is not observed in the averaged amplitudes of experimental *α* oscillations. Lastly, a further enhancement occurred when the noise amplitude was also increased (Fig. 5F) matching the experimental data (Fig. 5D).

To conclude, the present findings support the view that the deepening of anesthesia alters the E-I balance, changing the network toward increased excitation, thus amplifying the *α* oscillation amplitude. This effect can be recapitulated in our model by differentially decreasing the weight coefficient *ω*_1_ and *ω*_2_, showing that the model can reproduce changes in the event amplitude observed in vivo.

### *α* oscillation frequency reflects the diversity of thalamo-cortical activation

We next studied the role of the thalamus and cortex in the control of the frequency of the *α* oscillation. Our model suggested that the network connectivity is reflected in the *α* oscillation frequency (Fig. S4D-G) and that the frequency could vary with the amplitude similarly to the in vivo data (Fig. S5). First, from light to deep anesthesia, we observed an increase in the main frequency from a narrow peak at 5Hz to a continuous up to 16Hz (Fig. 6A-B, Fig. S2). More specifically, as shown in (Fig. 6C), the *α* oscillation frequency continuously increases with higher isoflurane dose (slope=7 e^−3^Hz.sec^−1^).

**Figure 6.**
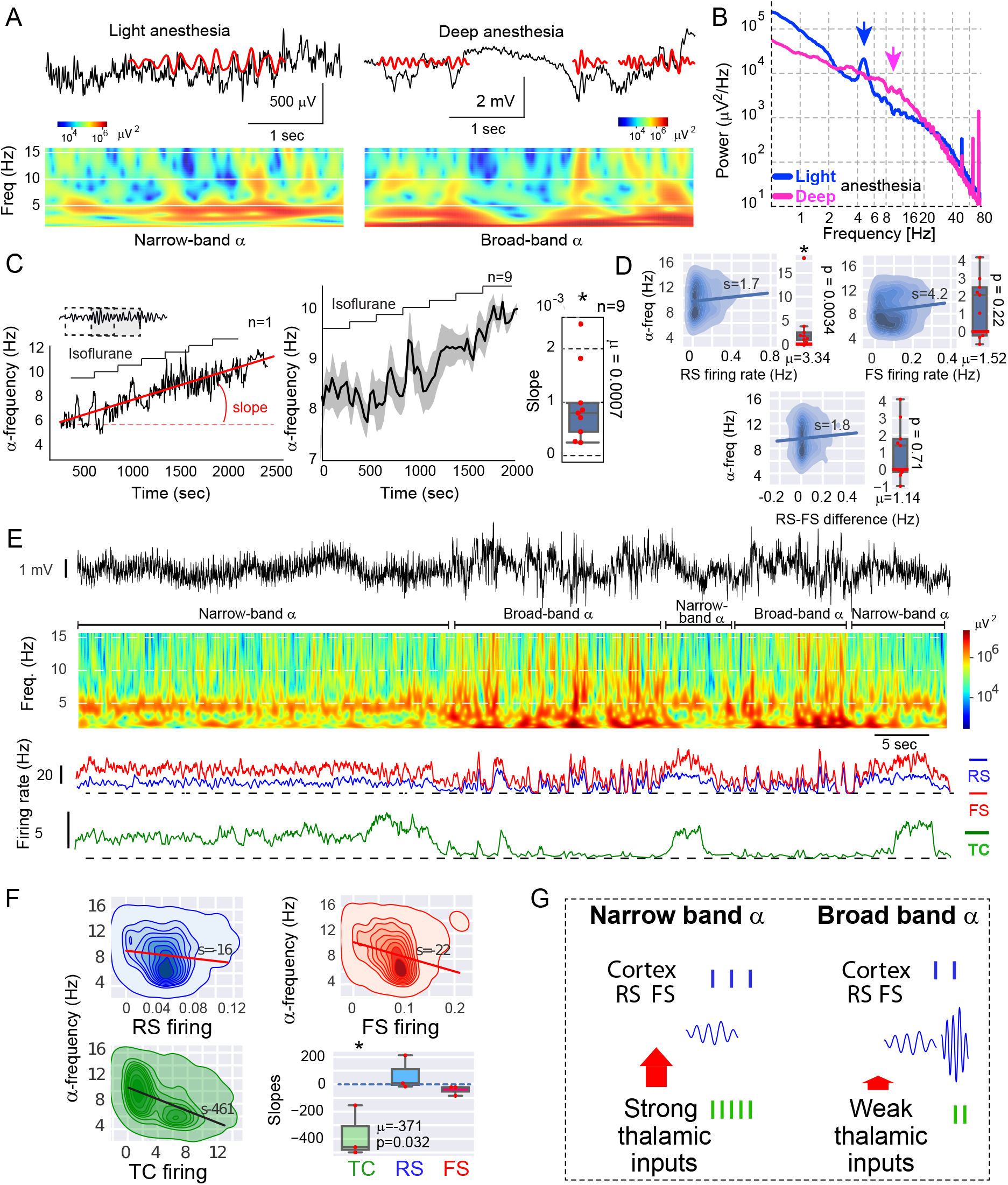
Increasing anesthesia depth redistributes *α* oscillation frequencies and uncouples thalamo-cortical inputs. **(A)** Example of raw LFPs (black), segmented *α* oscillations (red) and scalograms (color maps) under light (left) and deep (right) anesthesia. **(B)** LFP power spectra under light and deep anesthesia show different deviations (arrows) from the tendency either as single peak at 5Hz or as a bulge from 3 to 16Hz. **(C)** *α* oscillation frequency increases as the dose of isoflurane increases (same protocol as in Fig. 5A) in a single mouse (left) and at the population level (n=9 mice) (middle and right) (* *p <* 0.05 Wilcoxon test). **(D)** Density plot examples of *α* oscillation frequency vs. RS firing rate (top left), FS firing rate (top right) and RS-FS firing rate difference (bottom) and box plots for population (n=9 mice) distribution. **(E)** Top: Raw LFP (and coloured scalogram) with transition from broad-band *α* oscillations during deep anesthesia (with burst suppressions) to narrow-band *α* oscillations during light anesthesia. The isoflurane dose here is constant (0.7%) and the transitions from bust-suppression to *α* oscillations are spontaneous. Below: average firing rate of the cortical RS and FS populations and thalamic TC neurons during burst suppressions, broad-band frequency *α* oscillation and narrow-band frequency. Note the different thalamic activity between burst-suppression and narrow-band *α* oscillation **(F)** Left: density of the *α* oscillation frequency vs. neuron firing rate for different neuron types. Right : *α* oscillation frequency significantly correlates negatively with TC firing (n=3 mice). **(G)** Schematic diagram of *α* oscillation frequency modulation. High thalamic inputs (left) to the cortex result in narrow band frequency *α* oscillations whereas low thalamic inputs (right) induce broad-band frequency *α* oscillations.

To dissect the contribution of the cortical and thalamic network to this effect, we first correlated the firing rate of RS and FS neurons, and their difference (RS-FS) with the *α* oscillation frequency (Fig. 6D). There was a significant correlation between frequency and RS neuron firing, suggesting a decrease in frequency as anesthesia deepens. In addition, there was no correlation either with the FS firing rate or the difference between RS and FS firing rates (Fig. 6D). Therefore, the cortical activity does not play a direct role in the observed increase in *α* oscillation frequency at increasing anesthetic doses. Moreover, we found that a highly active thalamus is associated with a narrow frequency band of the *α* oscillation in the cortex, while a small thalamic input to the cortex leads to broadband *α* oscillation frequencies (Figs. 6E-F, Fig. S6). In addition, the TC neuron bursting activity associated with the increase of firing mostly consists of high-threshold spikes (i.e. low-frequency bursts [2]) and not the “classical” low-threshold spikes (i.e. high-frequency bursts)[28](Fig. S7).

In summary, a strong thalamic input imposes a stereotyped 5-Hz *α* oscillation frequency in the cortex, generated by RS-FS neuron interactions, while a low thalamic input leads to higher and broader *α* oscillation frequencies in the cortex (Fig. 6G). This suggests that the *α* oscillation frequency is most likely controlled by the reciprocal network connectivity, as predicted by our model.

### Coupled thalamo-cortical neuronal-mass models reproduce anesthesia dose-dependent frequency shifts

To better understand the increase in *α* oscillation frequency with increasing anesthetic dose, we use our modeling approach by coupling cortical and thalamic stochastic neural mass-models together (Fig. 7A) (see Methods). This allowed us to study the dynamics of *α* oscillations under large and low thalamocortical coupling modes as a proxy of the anesthetic doses (Fig. 7B). This system generates *α* oscillations at the dominant frequency of 5 Hz and 7 Hz when the coupling between cortex and thalamus is high and low, respectively. In addition, mixed *α* oscillations with both high and low frequencies (i.e broad-band) can emerge at intermediary doses (Fig. 7C). Indeed their proportion with respect to the strength of the coupling changes progressively. We observed a decreasing proportion of fast in parallel to an increasing proportion of slow *α* oscillations as the coupling strength increased (Fig. 7C). At the transition, we found the *α* oscillation with mixed-frequency (red curve in Fig. 7C). Interestingly, in this regime when fast, slow and mixed frequency *α* oscillations can occur alternatively (Fig. S8), studying the order of individual *α* spindle-like oscillation appearance showed that high-, mixed and low-frequency *α* oscillations appear in a random order and thus could reflect the broad-band *α* oscillation frequency of the *α* oscillation observed in the brain at high doses of anesthetics. Thus, the mass-model that accounts for thalamo-cortical interactions recapitulates the increase of *α* oscillation frequency at increasing anesthetic dose.

**Figure 7.**
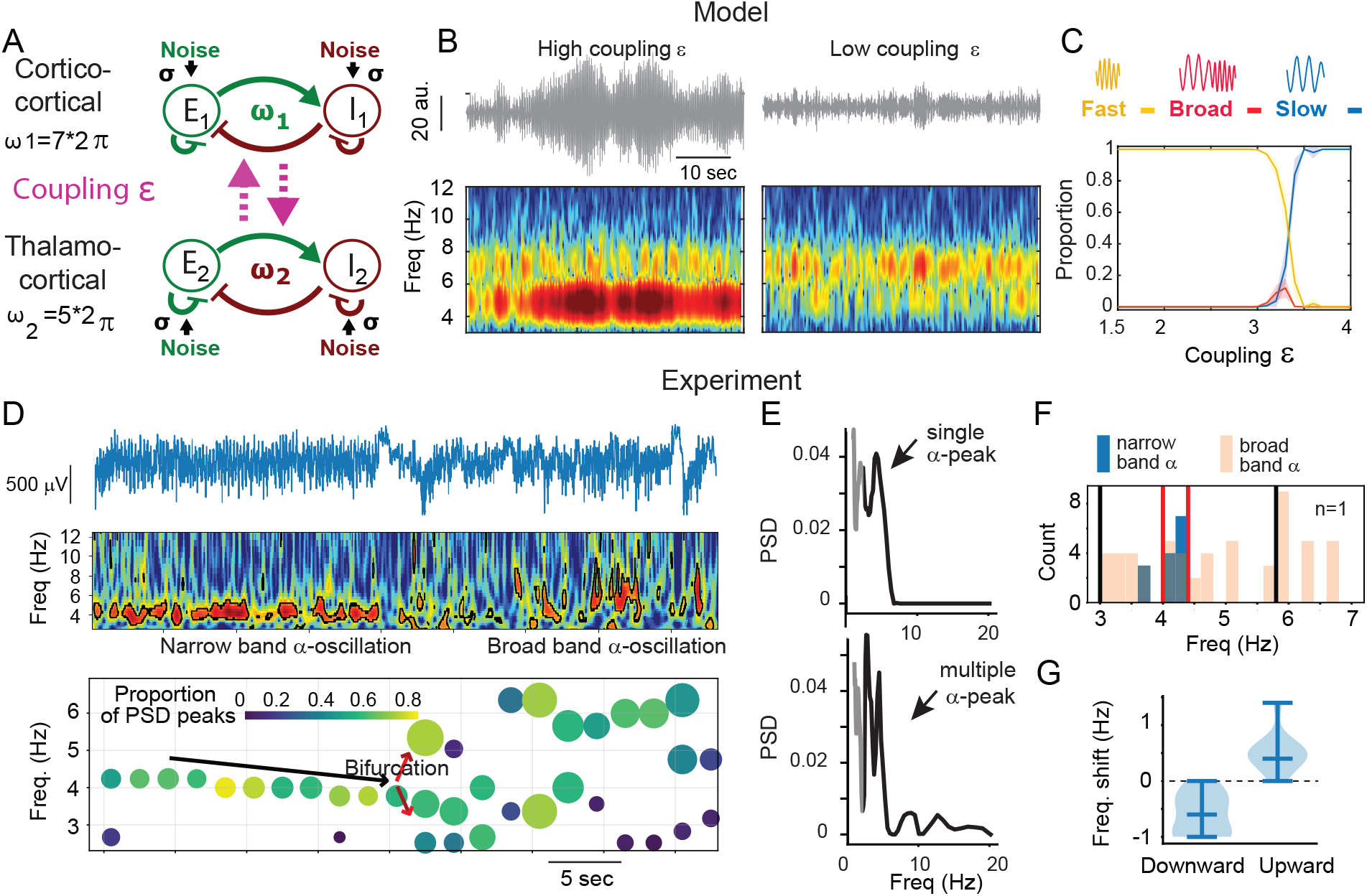
*α* oscillations of low and high frequencies reflect the coupling of two networks. **(A)** Schematic model of two coupled E-I networks (*ω*_1_=7*2*π, ω*_2_=5*2*π, λ*=1, *σ*=12. The coupling consists of *ϵ*. **(B)** Simulations with *ε*=3.1 (left) and *ϵ*=1.5 (right) and their relative LFP spectrograms. **(C)** Evolution of *α* oscillation-type proportions with respect to the coupling strength *ε* indicate a progressive increase of slow *α* oscillation compared to fast *α* oscillation when the coupling increases. **(D)** Top: LFP trace around the transition from narrow-band to broad-band mixed *α* oscillations. Middle: scalogram indicating time-frequency zones of maximal *α* oscillation intensity circled in black. Bottom: 4D-plot indicating for each 2-sec time bin the frequency of the power spectrum peaks (1,2 or 3 circles per time bin indicate multiple peaks), the proportion of the total Power Spectrum Density (PSD) (color coded) and the amplitude of the peak (size coded). Arrows indicate continuity during narrow-band *α* oscillations and a divergence during the mixed frequency broad-band *α* oscillations. **(E)** Power spectrum density of the masked LFP during the narrow band alpha (top) and of the LFP during the mixed frequency alpha (bottom). Only the peaks between 2.5 and 20Hz are considered (black vs. gray). **(F)** Distribution of the frequency peaks (from the example in (E)) before the bifurcation (blue) and after bifurcation (pink). Red and black lines indicate percentiles. **(G)** Averaged downward and upward frequency shift in Hz from the narrow-band median frequency (n=10 bifurcations) (n=4 mice).

Lastly, to characterize the transition from low 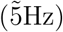 to broad-band *α* frequency, we selected spontaneous transition events (lasting 40 s) between these two frequency modes in the experimental data (Fig. 7D). As indicated in the transition depicted in the middle panel of Fig. 7D, the LFP scalogram shows a stable narrow-band at 5 Hz during the first 20 s, which is followed by a period characterized by broad-band *α* frequency. This transition (bifurcation between the two periods) is quantified in lower panel of Fig. 7D that is based on a time-frequency segmentation applied to the scalogram (see Methods). During the second period, multiple peaks are widely distributed around the narrow band *α* frequency (Fig. 7E-G, Fig. S9).

In conclusion, during anesthesia, the LFP can show transitions from a mono-frequency *α*-band to multiple bands, reflecting dynamic transitions, that are well reflected in our coupled-thalamo-cortical mass model.

## Discussion

The main findings of this study, combining in vivo LFP, single thalamic and cortical neuron recordings in mice with neuronal-mass modeling of *α* oscillations during general anesthesia show that: 1) the amplitude of the *α* oscillations reflects the depth of anesthesia with higher amplitude observed at increasing anesthetic dose, along with an increased E-I imbalance, 2) the frequency of the *α* oscillation bifurcates from a low narrow-band at low anesthetic doses to a continuum broad-band at higher doses during burst-suppression, 3) a neural-mass model consisting of two mutually interacting, excitatory and inhibitory neuronal populations, faithfully reproduces the features of the narrow-band *α* oscillations recorded in vivo. 4) two-interacting mass-models (i.e. thalamus and cortex) well reproduce the emergence of faster broad-band *α* oscillations that are observed at increasing anesthetic doses and suggest the causal role of a weaker thalamic input in the generation of these broad-band oscillations. Thus, the present framework offers a perspective on how *α* oscillations encode brain states that could be used to monitor the depth of anesthesia, potentially more accurately than the anesthetic dose.

### Study limitations

Before we examine the physiological and clinical implications of our results, we discuss the potential limitations of this study. First, differently from what is commonly observed in humans [1], the anesthetic-induced *α* oscillations in mice show a considerable amount of variability in frequency and in waxing and waning. This could be due to the different cortical influence between the two species and/or the stronger homeostasis and stability of the narrow-band *α* in humans. This issue, however, does not affect the main conclusions of our study.

Second, a preponderance of anesthetic-induced *α* oscillation activity is observed in humans over the frontal cortex. However, *α* oscillations at a lower power are observed in other human cortical regions, including the occipital cortex where our LFP and neuronal recordings were performed. The choice of occipital cortex was dictated by the technical limitations of inserting a probe horizontally into layer 4 of the frontal cortex in mice. Moreover, by recording in the occipital cortex we could also carefully confirm the position of the probe by measuring the known response of neurons to visual gratings.

### Determinants of *α* oscillations frequency across species, age and anesthesia depth

The dominant frequency of *α* oscillations recorded in EEG under anesthesia varies between mammalian species, as illustrated for mice and primates at 5Hz and 10Hz, respectively. They have the same frequency shift for these two conditions between species and are both associated with the spontaneous activity of the thalamocortical network [29, 26]. In fact, the 3-6 Hz frequency band in mice has been described as a potential evolutionary precursor of *α* oscillations [22] and *µ*-rhythms in rats [3].

The frequency of the *α* oscillations is also highly dependent on variables such as age, cognitive state and neurological conditions [30]. These differences could be related to the properties of the inherent neuronal circuit such as difference in cortical supra-granular layers [31] as they impact the specific resonance properties of the neocortex [32, 33] or the large difference of the thalamic pulvinar nuclei between species [34] that have been shown to be essential for the gating of *α* oscillations [29] or even connectivity differences between humans and mice [35, 36].

When the dose of the anesthetic increases, the *α* power is reduced, which could even lead to the loss of the *α* band (*α* suppression) [9]. Interestingly, as isoflurane increases, we found that the frequency of *α* oscillation bifurcates into two asymmetric branches: one characterized by low frequency and small power and the other by high frequencies, present mostly during burst-suppression (Fig. 7D). This slowdown is similar to those observed in human studies, where the frequency of *α* oscillation typically decreases with increasing doses of anesthetics [37].

Our modeling results suggest that the decrease in *α* oscillation frequency can be linked to the decrease of synaptic strength as it is observed in experimental conditions [38] whereas the increase in frequency associated with burst-suppression can be explained by changes in reciprocal thalamus-cortex coupling (Fig. 6G,7). In fact, when the cortical and thalamic networks are strongly coupled during light anesthesia, the *α* frequency is dominated by the narrow-band *α* oscillation whereas as anesthetic dose increases, the thalamic drive decreases, the cortical dynamics decouples, and higher broad-band frequency modes emerge in the cortex in *α* oscillations inter-spaced by isoelectric suppressions (Fig. 6). Thus, the increase in average frequency cannot be accounted for by local E-I cortical balance in the simple E-I 2-population model as it would predict a decrease in frequency. Instead, it reflects long-range thalamocortical interactions whose decrease removes some brakes on the cortical oscillators and allows higher *α* frequency activity. This interpretation is consistent with reports that anesthetics alter the excitability of TC cells and disrupt corticothalamic synchrony [39, 40]. It may also be possible that beyond the classic thalamocortical system generating the narrow-band low frequency *α* oscillation, some other E-I networks (independent of the thalamus) can affect cortical activity to generate *α* oscillations at a broad range of frequencies.

### *α* oscillation amplitude as a readout of E–I balance and firing variability

Our data show that *α* oscillations emerge amid similarly balanced firing of cortical excitatory RS and inhibitory FS neurons compared to activity in between *α* oscillation events (Fig. 1). However, the increase in amplitude across thousands of neurons was associated with shifted E-I ratios (Fig. 5C-D). Furthermore, increased variability in excitatory RS neuron firing also contributed to larger amplitude events (Fig. 5D-E), suggesting that both the mean firing rate and the temporal dispersion of the spiking are critical determinants of the size of the *α* oscillation envelope (Fig. 5F-G). When the thalamic inputs are strong (narrow-band low frequency *α*) during light anesthesia, the amplitude is highly regulated and relatively small compare to deeper anesthesia. These findings are consistent with the hypothesis that local thalamocortical and cortico-cortical interactions modulate the amplitude of the oscillation [2, 41]. Importantly, our neural-mass model captures this effect: the amplitude increased with increasing noise (*σ*) and with a shift in the effective E/I coupling. Thus, *α* oscillation amplitude can be interpreted as an emergent macroscopic readout of microscopic E-I network dynamics. This property could also extend to sleep states, which are particularly sensitive to perturbations of the homeostatic balance and can reveal latent hyper-excitability in vulnerable circuits, such as those observed in models of Alzheimer’s disease [42, 43]. These findings also suggest that amplitude fluctuations during low-arousal states may directly reflect homeostatic E-I regulation.

### Minimal population models of thalamo-cortical dynamics under anesthesia

Modeling thalamo-cortical rhythms in the 4 to 16 Hz range of activity serves to better characterize the mechanism of *α* oscillation genesis, but has faced two main challenges: 1) identifying the origins of the rhythmic activity in the thalamus and/or the cortex, and 2) characterizing the initiation and termination mechanisms. Rhythmic activity depends on specific membrane channels, receptors, and synapses [13, 14], while initiation and termination could depend on adaptive currents [44] or short-term synaptic plasticity [17]. One caveat of the most recent models is the presence of very synchronized assemblies, which are far from representing normal physiological conditions, and the high number of parameters which cannot be entirely justified as most of them are not measurable. Previous modeling studies [45, 16, 46, 47, 48] have also shown that synaptic dynamics, especially involving short-term plasticity, can produce transient oscillations during cortical Up-states, where noise induces sustained bursts through stochastic resonance, while others proposed to describe such transient activity, including mean-field or population-based neural field models [19, 49]. Instead, our model proposed a mechanism that keeps minimal biological constraints and assumptions (Figs. 3, 5, S4, S5), i.e. only includes excitatory and inhibitory populations, their synaptic connections, and some noise. In the second version of the model (Figs. 7, S7), the coupling of two similar networks to approximate the interactions of the thalamic and cortical circuits gives rise to *α* oscillations whose frequency depends on the coupling strength. The minimally constrained approach is flexible enough to be adapted to anesthetized mice and humans despite the large number of differences between these species.

### Leveraging the features of *α* oscillation for anesthesia monitoring and prediction

Although *α* oscillations appear as stochastic expressions of a random system, where individual events are unpredictable, their features (duration, amplitude, and time intervals) are tightly constrained by the model parameters. This paradigm shifts the emphasis from individual *α* oscillation events to their statistical evolution over time. The simplest form of our model of EEG *α* oscillation events (Fig. 3) could be used in real-time to monitor brain state stability by the continuous estimation of only 3 parameters (*λ, ω* and *σ*), thus providing a compact and physiologically interpretable representation of ongoing brain activity. In practice, this approach could enable the construction of a patient-specific numerical twin: a computational model that tracks the patient state stability based on the features of multiple *α* oscillation events computed over hundreds of seconds. This framework could also be used as a reference to quantify and predict the deviation at the level of a single *α* oscillation event.

Current EEG-based indices, such as the bispectral index (BIS), rely on heuristic signal features that are not designed to predict the arrival of deep sedation characterized by burst suppression or excessive sedation [9, 50, 10]. By contrast, tracking the features of *α* oscillation events and their underlying parameters may allow predicting transitions to deep sedation before these transitions occur [10]. In particular, inter-*α* oscillation event intervals can provide an early signal to predict the arrival of burst suppression [51, 52, 53].

### Anesthetic *α*-band activity reveal latent thalamo-cortical instabilities

Anesthesia could be used as a functional stress test to expose early neural circuit dysfunctions [43, 42]. In particular, in vulnerable states of neuro-degenerative diseases, such as mouse models of Alzheimer’s disease [54, 42, 43], anesthesia can challenge the E-I balance, even long before overt clinical symptoms appear. Indeed, while the healthy awake neocortex is dominated by inhibition [55], anesthesia reduces the balance between inhibition and excitation, thus creating particular conditions, which, in the case where hidden dyshomeostatic network properties preexist, can create altered oscillatory dynamics. Thus, the extraction of EEG *α* oscillation features during anesthesia could provide early predictions of the brain degenerative route. The rate of change in *α* frequency and amplitude under anesthesia may reflect anesthetic-induced thalamocortical decoupling, a framework providing a quantitative model that links excitability shifts to measurable features of *α* oscillations. Thus our modeling approach provides the formal tools to quantify such dysfunctions by linking neural circuit-level instabilities to *α* oscillation event characteristics.

In summary, we showed here that *α* oscillation events during anesthesia can be mechanistically random patterns, shaped by cortical E-I balance and thalamocortical coupling. Also, their changes in amplitude and frequency could be used to assess the degree of brain sedation, while changes of their features with time could offer predictive clinical information about brain state transitions.

## Material and Methods

### Neuropixels Recordings

C57BL/6J mice were bred in the local animal facility and maintained on a standard diet. Mouse data were collected according to the requirements of the local ethics committee (APAFIS 50287-2024070300528714, agreement ID D750512). 4-to 8-month-old mice, first underwent 1-% isoflurane anesthesia for cranial implantation. In the first two groups (n=4 and 9 mice respectively), a 384-channel Neuropixels 1.0 electrode was inserted into the primary visual cortex, parallel to layer 4 and its position was confirmed by histology and by the presence of receptive fields in the neuronal activity observed in response to white squares on a black background visual stimulation. In the third group (n=3 mice), the electrode was vertically inserted (AP-2.5mm, ML:2mm, DV: 3.8mm) so that it crosses the visual cortex and thalamus The first group was maintained at a steady level of anesthesia with 0.7% isoflurane. The second group underwent incremental anesthesia from 0.5% to 2.1% with 0.2% steps every 5 minutes (Fig.6). The anesthesia of the third group was constantly adjusted to capture the transitions from light to deep anesthesia. The recording procedures were similar to [56].

### Spike train and local field potential pre-processing

Statistical analysis involved the use of Phy [57] and Kilosort [58] in a semi-automatic procedure. Briefly, spikes extracted from the high-pass full-band (30 kHz) signal were clustered according to the spike waveform and spatial location of their best recording channel on the electrode [59]. Then based on well-characterized extracellular spike shapes of inhibitory neurons and excitatory neurons of the cortex and thalamus, spike waveforms with narrow spike shapes (trough-to-peak *<* 0.5*msec*) were classified as belonging to inhibitory cortical or thalamic neurons and those with broad spike shapes as excitatory cortical or thalamocortical neurons [60, 61].Thalamocortical axons, which reflect thalamic neuron activity impinging on layer 4 of the primary visual cortex, were identified based on the multi-phasic spike shape [56].

*α* oscillations in the local field potential were segmented from the filtered signal. The signal envelope amplitude was estimated from the distance from inferior to superior envelope limits (see EMD for the detection of the envelopes) of the band-pass -6 to 16 Hz-filtered signal (unless noted differently). The amplitude maxima above an upper threshold (3 standard deviations) define the presence of an *α* oscillation. The minimum value of the nearest-in-time envelope inferior to a lower threshold (1.5 x std) and anterior and posterior to the maximum is defined as the start and end of the *α* oscillation, respectively.

The variation in the envelope amplitude and *α* oscillation characteristics is only considered when they are outside suppressions detected based on the definition [9]. The local maxima of the histogram of each population firing are extracted over a sliding window of 100 seconds, and linearly fitted to extract the slope (Fig. 5C). Power spectra were estimated using multi-taper analysis [62] for each 5-min interval corresponding to a particular dose of anesthetics. The values are averaged over a sliding window of 0.6Hz. The power spectrum ratios were calculated on these windowed values. The variables were compared using the Wilcoxon sign rank test (paired samples) or the Mann-Whitney U test (unpaired samples), as appropriate. The significance level used was *α*=0.05.

### *α* oscillation segmentation using the Empirical Mode Decomposition

We use the empirical mode decomposition [63, 64] (EMD), defined for a signal *x* as follows:

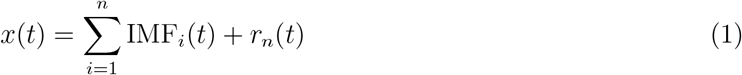

where IMF_*i*_(*t*) is the i-th intrinsic mode function and *r*_*n*_(*t*) is the residual after extracting *n* IMF [64, 65, 66]. In practice, we only use the first mode *n* = 1, allowing the identification of local extrema of *x*(*t*) from the upper envelope *u*(*t*) (resp. the lower envelope *l*(*t*)) (see Fig. 3A). We recall that *h*_1_(*t*) = *f* (*t*) − *m*(*t*), where 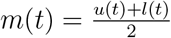

The segmentation of *α* oscillations is obtained between two consecutive minima with a single global maximum. To obtain *α* oscillation segmentation, we use a first threshold *T*_1_ = *σ* (standard deviation of the signal) to extract the beginning and end of the *α* oscillations at *T*_2_ = 3*σ* to extract the maximum amplitude, so that there is only one large maximum per *α* oscillation. The lower threshold *T*_1_ eliminates the possible fluctuation generated by the noise component of the amplitude. This procedure allows to segment the dynamics into *α* oscillations and periods in between, so that the max amplitude per *α* oscillation can be computed (Fig. 4B).

### E-I model spectral properties

The model equation is given by

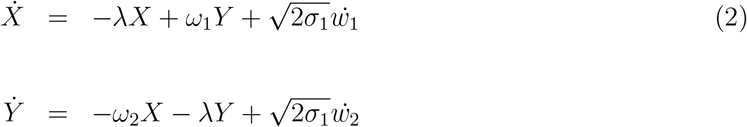

where ***w*** is the two-dimensional Brownian motion with mean zero. The oscillation is characterized by the two complex eigenvalues *µ*_*±*_ = −*λ ± iω*, where the main frequency is 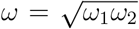. (The main frequency is *ω/*2*π*). We consider here the case where the noise amplitude *σ* is neither too small nor too large (of the order 1) and the ratio 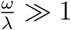, a situation that differs from fluctuations around a limit cycle oscillation, which is dominated by the deterministic component. Model 2 is a particular case of the Cowan-Wilson model [18] or can also result from the linearization of synaptic short-term plasticity mean-field model around the Up-state attractor [16, 67].

The steady-state distribution of the I-E dynamics is Gaussian [68, 69] in the variable ***s*** = (*x, y*)

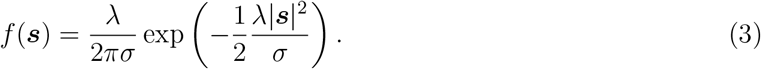

The distribution of *α* oscillation amplitudes is described by the equation for the radius 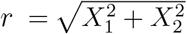

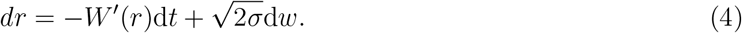

Where 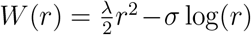. Thus, the noise amplitude has an equilibrium point at *W*^*′*^(*A*_*asymp*_) = 0 defined by 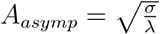. The distribution of *α* oscillation amplitude is solution of the associated Forward Fokker-Planck equation

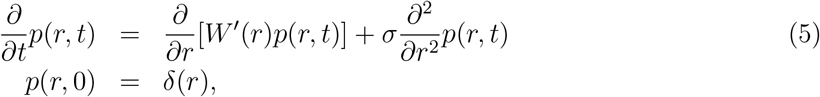

where δ is the Dirac δ− function. The stationary solution 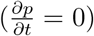 is given by

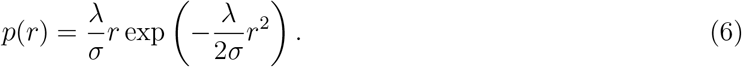

The power spectrum of the model equation is given by

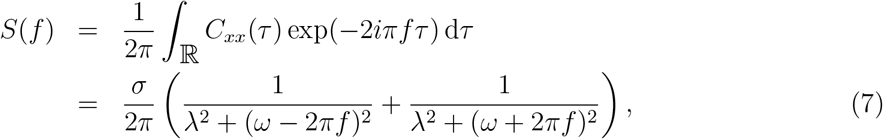

where *C*_*xx*_ is the autocorrelation function of the process first component [69]

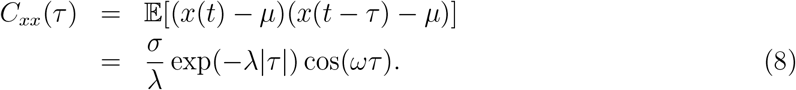

### Coupled *α* oscillation generators eq. 9

We propose here to couple two *α* oscillation generators that correspond to two stochastic focus model with prescribed resonance frequencies *ω*_1_ = 2*πf*_1_ and *ω*_2_ = 2*πf*_2_ . Specifically, the second system is coupled to the first by adding a small coupling parameter *ϵ* as follows:

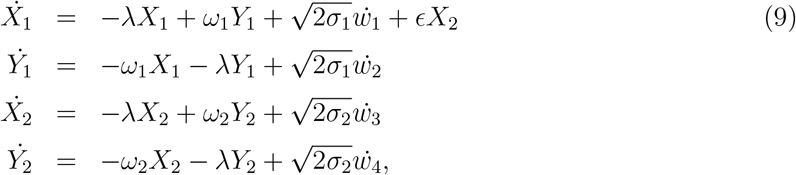

where *w*_*i*_ are Brownian motions and *λ* is the decaying parameter. The thalamocortical connectivity can be accounted for by several coupling parameters that we will introduce in the following system of equations where *X*_1_, *Y*_1_ can be associated to the cortex and *X*_2_, *Y*_2_ associated to the thalamus:

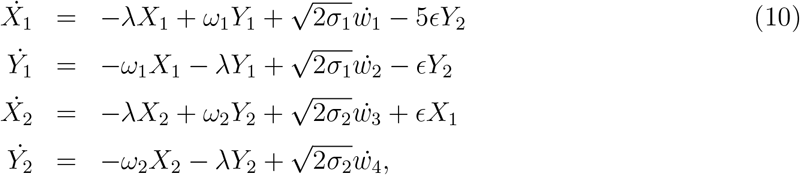

Specifically, to account for the thalamocortical projections on the cortex, through a long-range excitatory connection, we added a coupling term −*ϵY*_2_ on the second variable *Y*_1_ and a coupling term −5*ϵ* on the first variable *X*_1_. To account for the cortico-thalamic projection on the thalamus, via a long-range excitatory connection, we added a coupling term −*ϵX*_1_. The one hand from thalamocortical neuron axons projecting to inhibitory and excitatory neuron mostly from layer 4 of the cortex and 2) on the other hand from cortico-thalamic axons projecting from layer six pyramidal neurons to inhibitory reticular thalamic and thalamic neurons.

### Measuring slow, fast and intermediate *α* oscillation sequence randomness using Wald Wolfowitz and Entropy

To evaluate the randomness of the three-valued *α* oscillation sequence (*S, F, M*), we use the Wald-Wolfowitz run test [70, 71]. A run is defined as a consecutive sequence of identical values. The null hypothesis *H*_0_ assumes that the sequence is random. The number of runs under H0 can be approximated by the expected number of runs given by the following expressions

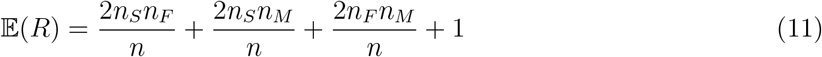

where *n*_*S*_, *n*_*F*_, and *n*_*M*_ are the counts of slow, fast, and mixed *α* oscillations, and *n* = *n*_*A*_ + *n*_*B*_ + *n*_*C*_ is the total number of *α* oscillations. The variance of the number of runs is calculated as

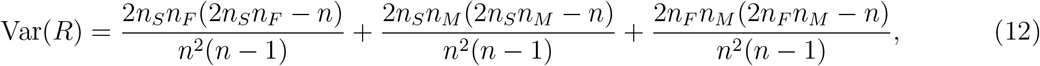

and the test statistic *Z* is

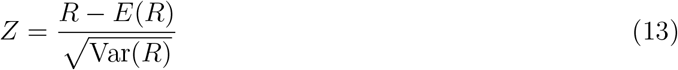

where *R* is the observed number of runs. The value of *Z* is compared to critical values from the standard normal distribution. If |*Z*| exceeds the critical value 1.96 for *α* = 0.05, H0 is rejected, indicating that the sequence is not random.

### Bifurcation analysis on spectrogram

We describe here a method for segmenting *α* oscillations that could overlap in time, but can be resolved in the time-frequency domain, as they separate in their frequency (Fig. 7D). The method starts with building the scalogram with the Morlet wavelet. We then segmented the image based on thresholding the upper 75^*th*^-percentile of the scalogram heatmap. The result provides clusters of high power *α* oscillations. In a second step, we further segment the long *α* oscillations, characterized by a number of pixels larger than the threshold *T* = 20 corresponding to the median value. This procedure provides a mask (only 0 or 1) for the time-frequency *α* oscillation segmentation (black contour Fig.7D). To extract the *α* oscillations, we apply a sliding window of 5 s with 3 s overlap to get a time-bin histogram of the segmented frequency *F*_*t*_(*f*), which we normalize by the total integral 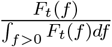 (Fig. 7E). Then, on each histogram, we detect each prominence peak above the arbitrary threshold 0.1 and their individual width (Fig. 7D bottom).

Note that the larger the circle radius is, the wider is the peak.

We further develop an algorithm to segment the emergence of several *α* oscillation branches in the time-frequency domain. The scheme is described (Fig. S9) as follows:

1. Compute the distributions of the circle frequencies before and after a possible bifurcation.
2. Extract the proportion *p*_*before*_ of circles within a single narrow band frequency range [*f*_0_, *f*_1_].
3. Compute the proportion *p*_*after*_ in the same frequency range [*f*_0_, *f*_1_] as in the previous step, but now after a bifurcation has occurred (this proportion should be lower compared to the one computed before the bifurcation).
4. Identify the left and right bands to account for the same total power proportion as the band before bifurcation. To do so, we compute the upper frequency limit *f*_*r*_ so that the proportion in the band [*f, f* ] plus *P* is equal to 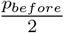. We repeated a similar procedure to estimate the lower band *f*_*l*_.

This procedure allows identifying two regions that contain each *α* oscillation branch after the bifurcation.

## Author contributions

Conceptualization, F.D.,J.S.,D.H.,V.C, N.R.; methodology, F.D., C.S, PO.M, J.S., D.H., Software, F.D., C.S. PO.M.,J.S.,D.H. Writing, F.D., V.C, D.H.”

## Supplemental information

Document S1. Figures S1–S9 and supplemental references

## Resource availability

Data and code can be provided upon request to the corresponding authors.

## Acknowledgment

D.H. research is supported by an ANR-23-CE46-0016 AnalysisSpectralEEG and AstroXcite and the European Research Council (ERC) under the European Union’s Horizon 2020 research and innovation program (grant agreement No 882673) and PoC 101212658.

## References

[1] Purdon, P. L. et al. Electroencephalogram signatures of loss and recovery of consciousness from propofol. Proceedings of the National Academy of Sciences 110, E1142–E1151 (2013).

[2] Hughes, S. W. et al. Synchronized oscillations at α and θ frequencies in the lateral geniculate nucleus. Neuron 42, 253–268 (2004).

[3] Hughes, S. W. & Crunelli, V. Thalamic mechanisms of eeg alpha rhythms and their pathological implications. The Neuroscientist 11, 357–372 (2005).

[4] Traub, R. D. et al. Layer 4 pyramidal neuron dendritic bursting underlies a post-stimulus visual cortical alpha rhythm. Communications biology 3, 230 (2020).

[5] Başar, E. A review of alpha activity in integrative brain function: fundamental physiology, sensory coding, cognition and pathology. International Journal of Psychophysiology 86, 1–24 (2012).

[6] Cimenser, A. et al. Tracking brain states under general anesthesia by using global coherence analysis. Proceedings of the National Academy of Sciences 108, 8832–8837 (2011).

[7] Altwegg-Boussac, T. et al. Cortical neurons and networks are dormant but fully responsive during isoelectric brain state. Brain 140, 2381–2398 (2017).

[8] Yin, M. et al. Synchronicity of pyramidal neurones in the neocortex dominates isoflurane-induced burst suppression in mice. British Journal of Anaesthesia 134, 1122–1133 (2025).

[9] Cartailler, J., Parutto, P., Touchard, C., Vallée, F. & Holcman, D. Alpha rhythm collapse predicts iso-electric suppressions during anesthesia. Communications biology 2, 1–10 (2019).

[10] Sun, C. & Holcman, D. Combining transient statistical markers from the eeg signal to predict brain sensitivity to general anesthesia. Biomedical Signal Processing and Control 77, 103713 (2022).

[11] Sleigh, J., Voss, L., Steyn-Ross, M., Steyn-Ross, D. & Wilson, M. Modelling sleep and general anaesthesia. In Sleep and anesthesia: neural correlates in theory and experiment, 21–41 (Springer, 2011).

[12] Bhattacharya, B. S., Coyle, D. & Maguire, L. P. A thalamo–cortico–thalamic neural mass model to study alpha rhythms in alzheimer’s disease. Neural networks 24, 631–645 (2011).

[13] Ching, S., Purdon, P. L., Vijayan, S., Kopell, N. J. & Brown, E. N. A neurophysiological– metabolic model for burst suppression. Proceedings of the National Academy of Sciences 109, 3095–3100 (2012).

[14] Soplata, A. E. et al. Rapid thalamocortical network switching mediated by cortical synchronization underlies propofol-induced eeg signatures: a biophysical model. Journal of Neurophysiology 130, 86–103 (2023).

[15] Bastiaens, S. P., Momi, D. & Griffiths, J. D. A comprehensive investigation of intracortical and corticothalamic models of the alpha rhythm. PLOS Computational Biology 21, e1012926 (2025).

[16] Holcman, D. & Tsodyks, M. The emergence of up and down states in cortical networks. PLoS computational biology 2, e23 (2006).

[17] Zonca, L. & Holcman, D. Emergence and fragmentation of the alpha-band driven by neuronal network dynamics. PLoS Computational Biology 17, e1009639 (2021).

[18] Wilson, H. R. & Cowan, J. D. A mathematical theory of the functional dynamics of cortical and thalamic nervous tissue. Kybernetik 13, 55–80 (1973).

[19] Brunel, N. Dynamics of sparsely connected networks of excitatory and inhibitory spiking neurons. Journal of computational neuroscience 8, 183–208 (2000).

[20] Buzsaki, G. Rhythms of the Brain (Oxford University Press, 2006).

[21] Dora, M., Jaffard, S. & Holcman, D. The wqn algorithm for eeg artifact removal in the absence of scale invariance. IEEE Transactions on Signal Processing (2024).

[22] Senzai, Y., Fernandez-Ruiz, A. & Buzsáki, G. Layer-specific physiological features and interlaminar interactions in the primary visual cortex of the mouse. Neuron 101, 500–513 (2019).

[23] Loison, V. et al. Mapping general anesthesia states based on electro-encephalogram transition phases. NeuroImage 285, 120498 (2024).

[24] You, Y. et al. Anesthetic spindles serve as eeg markers of the depth variations in anesthesia induced by multifarious general anesthetics in mouse experiments. Frontiers in Pharmacology 15, 1474923 (2024).

[25] Steriade, M. & Deschenes, M. The thalamus as a neuronal oscillator. Brain Research Reviews 8, 1–63 (1984).

[26] Nestvogel, D. B. & McCormick, D. A. Visual thalamocortical mechanisms of waking statedependent activity and alpha oscillations. Neuron 110, 120–138 (2022).

[27] Dayan, P. & Abbott, L. F. Theoretical neuroscience: computational and mathematical modeling of neural systems (MIT press, 2005).

[28] Llinás, R. & Jahnsen, H. Electrophysiology of mammalian thalamic neurones in vitro. Nature 297, 406–408 (1982).

[29] Saalmann, Y. B., Pinsk, M. A., Wang, L., Li, X. & Kastner, S. The pulvinar regulates information transmission between cortical areas based on attention demands. science 337, 753–756 (2012).

[30] Mierau, A., Klimesch, W. & Lefebvre, J. State-dependent alpha peak frequency shifts: Experimental evidence, potential mechanisms and functional implications. neuroscience 360, 146–154 (2017).

[31] Kalmbach, B. E. et al. h-channels contribute to divergent intrinsic membrane properties of supragranular pyramidal neurons in human versus mouse cerebral cortex. Neuron 100, 1194–1208 (2018).

[32] Halgren, M. et al. The generation and propagation of the human alpha rhythm. Proceedings of the National Academy of Sciences 116, 23772–23782 (2019).

[33] David, F., Borel, M., Ayub, S., Ruther, P. & Gentet, L. J. Layer-specific stimulations of parvalbumin-positive cortical interneurons in mice entrain brain rhythms to different frequencies. Cerebral Cortex 33, 8286–8299 (2023).

[34] Cortes, N., Ladret, H. J., Abbas-Farishta, R. & Casanova, C. The pulvinar as a hub of visual processing and cortical integration. Trends in Neurosciences 47, 120–134 (2024).

[35] Campagnola, L. et al. Local connectivity and synaptic dynamics in mouse and human neocortex. Science 375, eabj5861 (2022).

[36] Hunt, S. et al. Strong and reliable synaptic communication between pyramidal neurons in adult human cerebral cortex. Cerebral Cortex 33, 2857–2878 (2023).

[37] Hight, D., Voss, L. J., Garcia, P. S. & Sleigh, J. Changes in alpha frequency and power of the electroencephalogram during volatile-based general anesthesia. Frontiers in Systems Neuroscience 11, 36 (2017).

[38] Vizuete, J. A., Pillay, S., Diba, K., Ropella, K. M. & Hudetz, A. G. Monosynaptic functional connectivity in cerebral cortex during wakefulness and under graded levels of anesthesia. Frontiers in Integrative Neuroscience 6, 90 (2012).

[39] Ching, S., Cimenser, A., Purdon, P. L., Brown, E. N. & Kopell, N. J. Thalamocortical model for a propofol-induced α-rhythm associated with loss of consciousness. Proceedings of the National Academy of Sciences 107, 22665–22670 (2010).

[40] Brown, E. N., Purdon, P. L. & Van Dort, C. J. General anesthesia and altered states of arousal: a systems neuroscience analysis. Annual review of neuroscience 34, 601–628 (2011).

[41] Veit, J., Handy, G., Mossing, D. P., Doiron, B. & Adesnik, H. Cortical vip neurons locally control the gain but globally control the coherence of gamma band rhythms. Neuron 111, 405–417 (2023).

[42] Zarhin, D. et al. Disrupted neural correlates of anesthesia and sleep reveal early circuit dysfunctions in alzheimer models. Cell Reports 38, 110268 (2022).

[43] Slutsky, I. Linking activity dyshomeostasis and sleep disturbances in alzheimer disease. Nature Reviews Neuroscience 25, 272–284 (2024).

[44] Lüthi, A. & McCormick, D. A. Periodicity of thalamic synchronized oscillations: the role of ca2+-mediated upregulation of ih. Neuron 20, 553–563 (1998).

[45] Destexhe, A. Modelling corticothalamic feedback and the gating of the thalamus by the cerebral cortex. Journal of Physiology-Paris 94, 391–410 (2000).

[46] Verechtchaguina, T., Sokolov, I. M. & Schimansky-Geier, L. First passage time densities in resonate-and-fire models. Physical Review E—Statistical, Nonlinear, and Soft Matter Physics 73, 031108 (2006).

[47] Duc, K. D., Schuss, Z. & Holcman, D. Oscillatory decay of the survival probability of activated diffusion across a limit cycle. Physical Review E 89, 030101 (2014).

[48] Dao Duc, K. et al. Synaptic dynamics and neuronal network connectivity are reflected in the distribution of times in up states. Frontiers in Computational Neuroscience 9, 96 (2015).

[49] Brunel, N. & Wang, X.-J. What determines the frequency of fast network oscillations with irregular neural discharges? i. synaptic dynamics and excitation-inhibition balance. Journal of neurophysiology 90, 415–430 (2003).

[50] Shao, Y. R. et al. Low frontal alpha power is associated with the propensity for burst suppression: An electroencephalogram phenotype for a “vulnerable brain”. Anesthesia and analgesia 131, 1529 (2020).

[51] Chemali, J., Ching, S., Purdon, P. L., Solt, K. & Brown, E. N. Burst suppression probability algorithms: state-space methods for tracking eeg burst suppression. Journal of neural engineering 10, 056017 (2013).

[52] Plummer, G. S. et al. Electroencephalogram dynamics during general anesthesia predict the later incidence and duration of burst-suppression during cardiopulmonary bypass. Clinical Neurophysiology 130, 55–60 (2019).

[53] Sleigh, J. W., Scheib, C. M. & Sanders, R. D. General anaesthesia and electroencephalographic spindles. Trends in Anaesthesia and Critical Care 1, 263–269 (2011).

[54] Shoob, S. et al. Deep brain stimulation of thalamic nucleus reuniens promotes neuronal and cognitive resilience in an alzheimer’s disease mouse model. Nature Communications 14, 7002 (2023).

[55] Haider, B., Häusser, M. & Carandini, M. Inhibition dominates sensory responses in the awake cortex. Nature 493, 97–100 (2013).

[56] Sibille, J., Gehr, C. & Kremkow, J. Efficient mapping of the thalamocortical monosynaptic connectivity in vivo by tangential insertions of high-density electrodes in the cortex. Proceedings of the National Academy of Sciences 121, e2313048121 (2024).

[57] Rossant, C. et al. Spike sorting for large, dense electrode arrays. Nature neuroscience 19, 634–641 (2016).

[58] Pachitariu, M., Steinmetz, N. A., Kadir, S. N., Carandini, M. & Harris, K. D. Fast and accurate spike sorting of high-channel count probes with kilosort. Advances in neural information processing systems 29 (2016).

[59] Pachitariu, M., Sridhar, S., Pennington, J. & Stringer, C. Spike sorting with kilosort4. Nature methods 21, 914–921 (2024).

[60] Barthó, P. et al. Characterization of neocortical principal cells and interneurons by network interactions and extracellular features. Journal of neurophysiology 92, 600–608 (2004).

[61] McCafferty, C. et al. Cortical drive and thalamic feed-forward inhibition control thalamic output synchrony during absence seizures. Nature neuroscience 21, 744–756 (2018).

[62] Gorgolewski, K. J. et al. Nipype: a flexible, lightweight and extensible neuroimaging data processing framework in Python. 0.12.0-rc1 (2016). URL 10.5281/zenodo.50186.

[63] Jaffard, S., Meyer, Y. & Ryan, R. D. Wavelets: tools for science and technology (SIAM, 2001).

[64] Flandrin, P., Rilling, G. & Goncalves, P. Empirical mode decomposition as a filter bank. IEEE signal processing letters 11, 112–114 (2004).

[65] Wu, Z. & Huang, N. E. Ensemble empirical mode decomposition: A noise-assisted data analysis method. Advances in Adaptive Data Analysis 1, 1–41 (2009).

[66] Wu, H.-T. et al. An integrated complete ensemble empirical mode decomposition with adaptive noise to optimize lstm for significant wave height forecasting. Journal of Marine Science and Engineering 9, 631–645 (2021).

[67] Zonca, L., Dossi, E., Rouach, N. & Holcman, D. Computational methods and algorithms to segment and model recurrent bursting events in long-time series springer nature book about neuromethods. In New Aspects in Analyzing the Synaptic Organization of the Brain, 323–370 (Springer, 2024).

[68] Risken, H. Fokker-planck equation. In The Fokker-Planck Equation, 63–95 (Springer, 1996).

[69] Schuss, Z. Theory and applications of stochastic processes: an analytical approach, vol. 170 (Springer Science & Business Media, 2009).

[70] Bradley, J. V. Distribution-free statistical tests, vol. 60 (United States Air Force, 1960).

[71] Sheskin, D. J. Handbook of parametric and nonparametric statistical procedures (Chapman and hall/CRC, 2003).

