## Supplementary material for "Cortico-thalamic excitatory-inhibitory dynamics generates *α*-wave transient patterns encoding anesthetic brain states"

February 20, 2026

### 1 Supplementary Figures

#### Figure S1

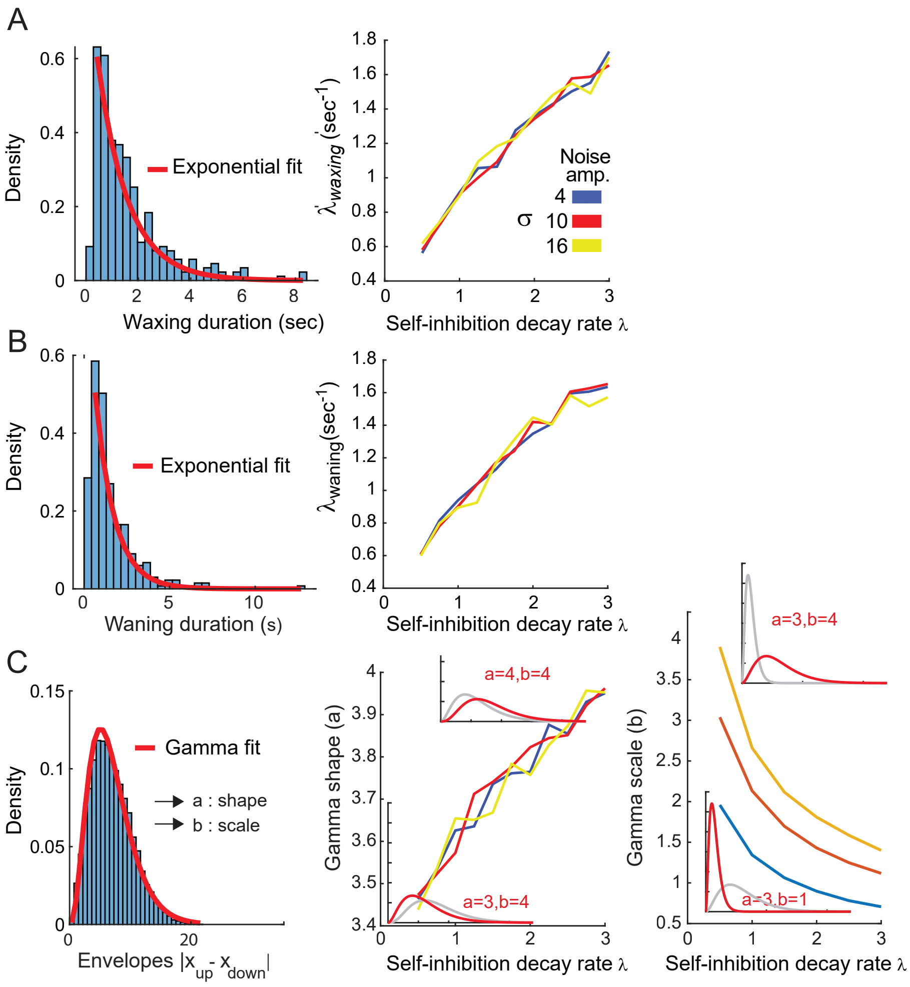


**Figure S1: Characteristics of simulated $\alpha$ oscillations vary with the self-inhibition decay rate and noise level (A)** Left: distribution of the duration of the $\alpha$ waxing for the same parameters of the model simulations shown in Fig. 4C-E. Red line indicates the fit with an exponential function. Right: the decay rate $\lambda_{waxing}$ varies linearly with the self-inhibition decay rate $\lambda$ but is invariant to noise amplitude $\sigma$ changes (color-coded). **(B)** Same as in A but for the duration of the $\alpha$ waning. Note the similarities with the waxing in A. **(C)** Left: distribution of the filtered signal envelope is well fitted with a Gamma distribution $\frac{1}{\Gamma(a)\,b^a}\,x^{a-1}e^{-x/b}$, (red line) $\Gamma(a)$ is the gamma function $\Gamma(a) = \int_0^\infty t^{a-1}e^{-t}\,dt$. Middle and Right: The two parameters indicating the shape (a) and the scale (b) vary distinctively with the process self-inhibitory decay parameter ($\lambda$) and the noise amplitude (colour-coded as in A). For middle and right plot, respectively, two gamma functions are represented illustrating two extrema values (red vs grey) for the shape (a) and the scale (b) parameters, respectively. Overall the increase of the self-inhibition decay rate tends to move the distribution toward small envelope amplitudes (i.e. small ($b$) values) and more symmetric (i.e gaussian shape instead of an exponentially decaying shape) values around the mean (i.e. high ($a$) values).

### Figure S2



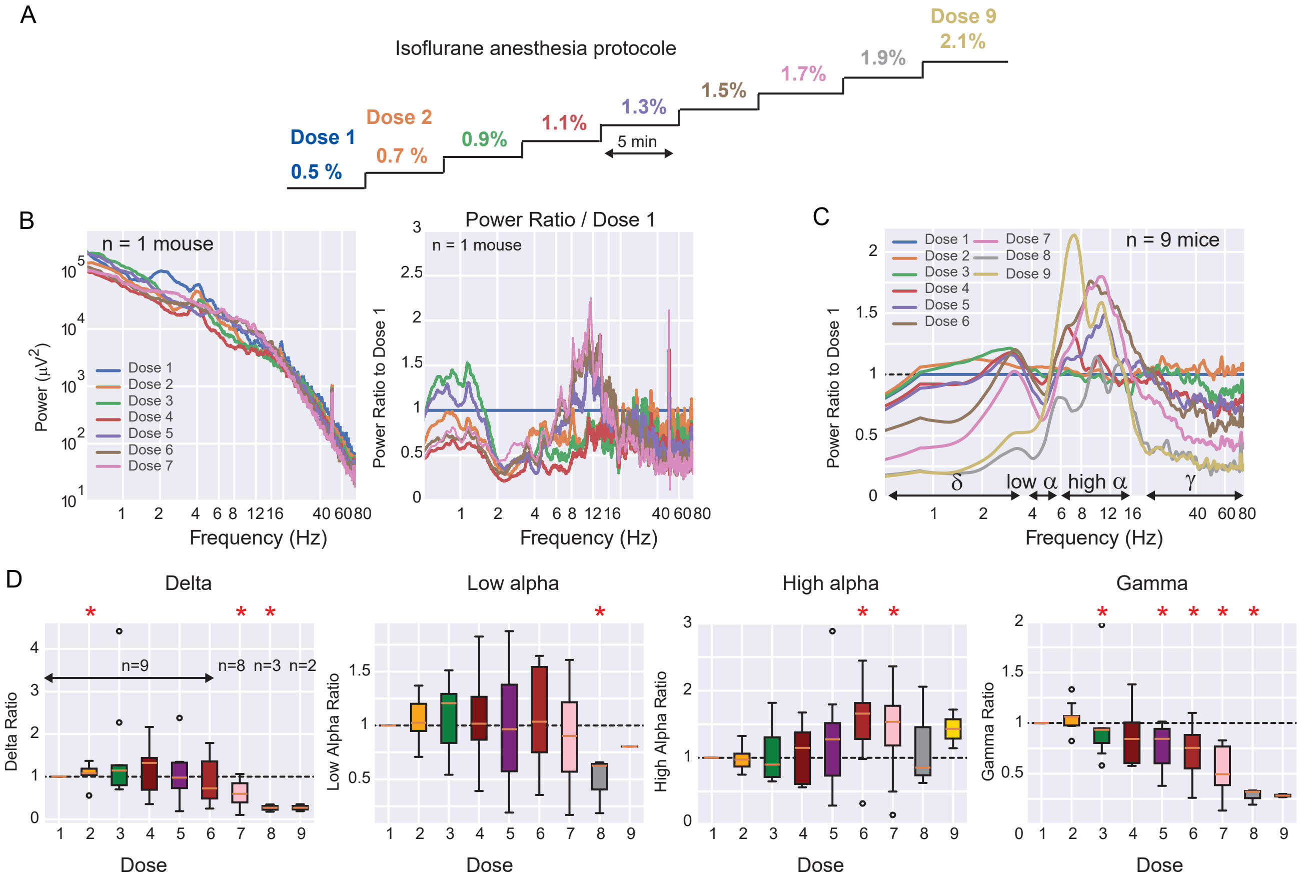


**Figure S2: Increase of the anesthetic dose enhances $\alpha$ band activity.** **(A)** Schematic drawing of the experimental protocol: the anesthetic dose starts from 0.5%, increases by 0.2% points every 5 min until 1.7% (n=9 mice). **(B)** Left: LFP power spectrum for 7 incremental doses of the anesthetic. Right: LFP ratio relative to the lowest dose (0.5%). **(C)** Average modulation for the power spectrum (from n=7 doses, n=9 mice to 9 doses, n=2 mice). The curves are smoothed for the representation. Power modulation affects the $\delta$ (0.5-3 Hz), low (4-6 Hz) and high (6-16 Hz) $\alpha$ and $\gamma$ (20-80 Hz) frequency bands. **(D)** Relative change of power (ratio compared to the minimal dose) in the $\delta$, low-$\alpha$ (4-5 Hz) high-$\alpha$ (6-16 Hz) and $\gamma$ bands. Only the high-$\alpha$ band shows an increase up to the highest doses, whereas only the $\gamma$ shows a constant decrease up to the highest doses of anesthetic. $*$ : p¡0.05 (Mann Whitney U test, n indicates the number of animals). $\circ$ marks outliers, defined as points lying more than 1.5x the interquartile range ($IQR = Q_3 - Q_1$) from the box. In the box plots, the yellow line is the median, the box edges are $Q_1$ and $Q_3$ and the whiskers extend to the most extreme non-outlier values.

### Figure S3

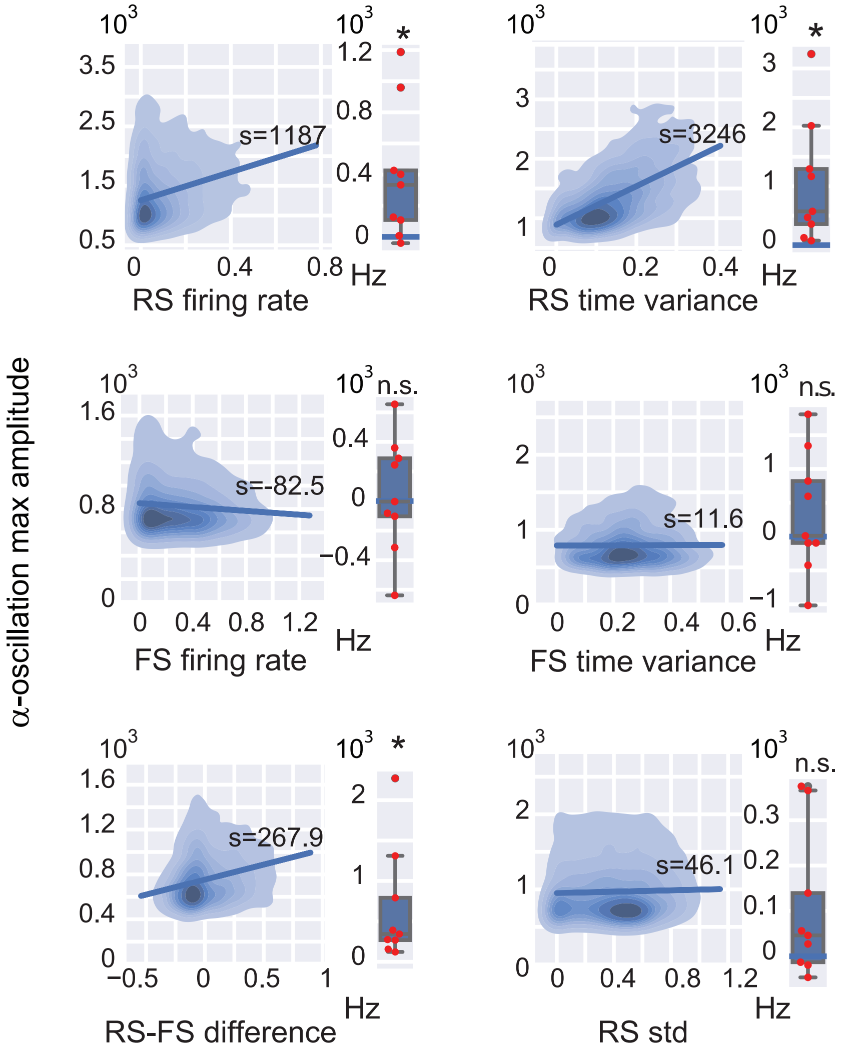


**Figure S3: Neuronal correlates of $\alpha$-oscillation amplitude.** Left column: from top, density plots of $\alpha$ oscillation maximal amplitude vs. RS firing rate, FS firing rates and RS–FS firing rate difference for 1 mouse. Right column: from top, same density plots of $\alpha$ oscillation maxima vs. RS firing time variance computed between 0.5 s and $\alpha$ oscillation maxima, same for FS firing time variance and firing variance across RS neurons. The slope S of the linear regression is indicated for each density plot. Box plots on the right of each diagram indicate the distribution of slopes ($*p < 0.05$, Wilcoxon test, n=9 mice).

### Figure S4

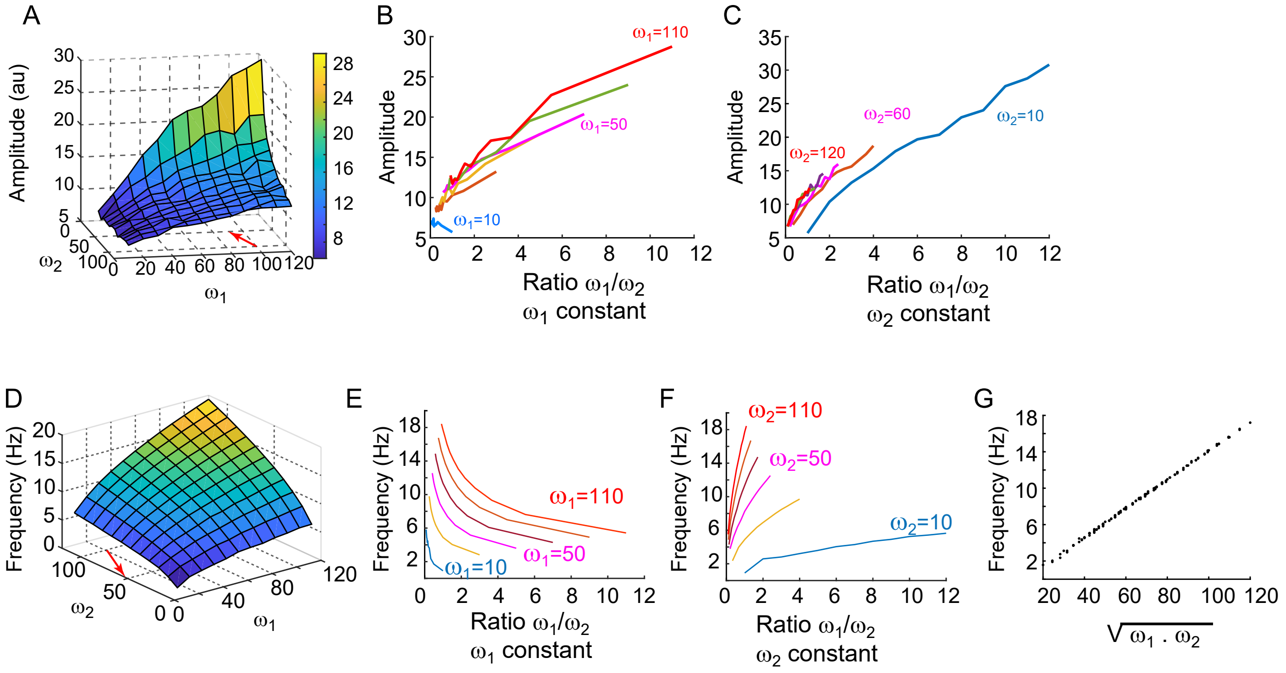


**Figure S4: $\alpha$ oscillation amplitude and frequency variations with $\omega_1$ and $\omega_2$ (A)** $\alpha$ oscillation maximal amplitudes (color code) varies as the excitatory and inhibitory connections $\omega_1$ and $\omega_2$ change. The amplitude strongly increases with $\omega_1$ and slightly decrease with $\omega_2$. The red arrow qualitatively indicates the hypothesized effect of an increase of anesthetic dose on the excitatory and inhibitory components of the model i.e. $\omega_1$ and $\omega_2$ (in the $\omega_1 * \omega_2$ plane) based on the experimental observation in Fig. 5C. i.e. both excitatory and inhibitory neuronal activity decrease and the inhibitory one decrease more than the excitatory one. The movement in the $\omega_1 * \omega_2$ plane result in an increase in amplitude (vertical axis). **(B)** The amplitude increases with $\omega_1/\omega_2$ while $_1$ is held constant. **(C)** The amplitude also increases with $\omega_1/\omega_2$ while $\omega_2$ is held constant. **(D)** The $\alpha$ oscillation frequency (color code) increases as the excitatory and inhibitory connection $\omega_1$ and $\omega_2$ increase. The red arrow, like in (A) indicates the hypothesized effect of isoflurane on the excitatory ($\omega_1$) and inhibitory ($\omega_2$) components. Here the model predict a decrease of frequency (vertical axis). **(E)** The $\alpha$ oscillation frequency decreases as the ratio $\omega_1/\omega_2$ increases ($\omega_1$ being held constant) **(F)** The $\alpha$ frequency increases as the ratio $\omega_1/\omega_2$ increases ($\omega_2$ being held constant). **(G)** The $\alpha$ frequency linearly increases with $\sqrt{\omega_1 . \omega_2}$.

### Figure S5



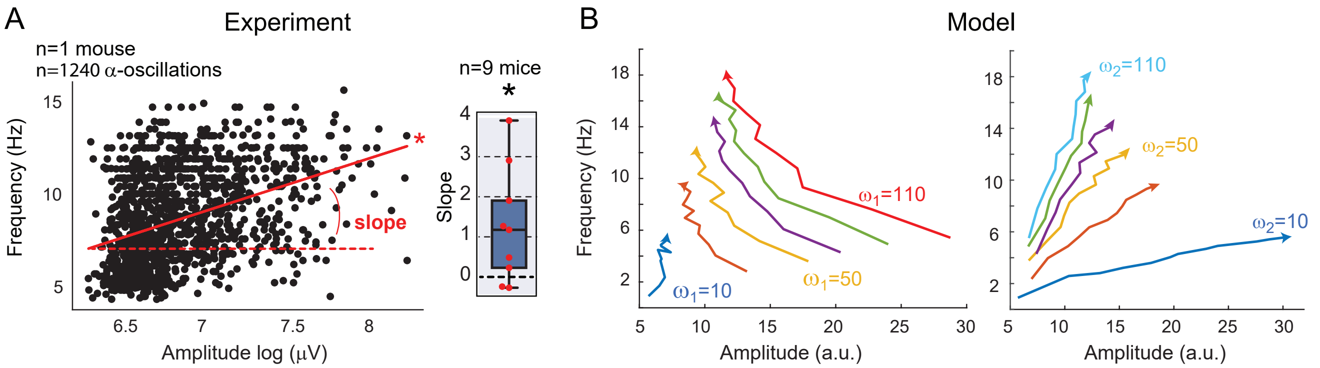


**Figure S5: Frequency-amplitude correlation for experimental and simulated $\alpha$ oscillations. (A)** Left: example of $\alpha$ oscillation frequency vs amplitude (n=1 mouse). Right: the distribution of linear regression factors (slope) is significantly positive (p=0.008, n=9 mice). **(B)** Frequency-amplitude relationship for $\omega_1$ and $\omega_2$ varying between 10 and 120 (left for $\omega_1$ constant; right for $\omega_2$ constant). The arrows on the curves indicate the increase of $\omega_2$ on the left and of $\omega_1$ on the right. This shows that the model cannot explain the amplitude frequency positive correlation observed experimentally when both $\omega_1$ and $\omega_2$ decrease.

### Figure S6



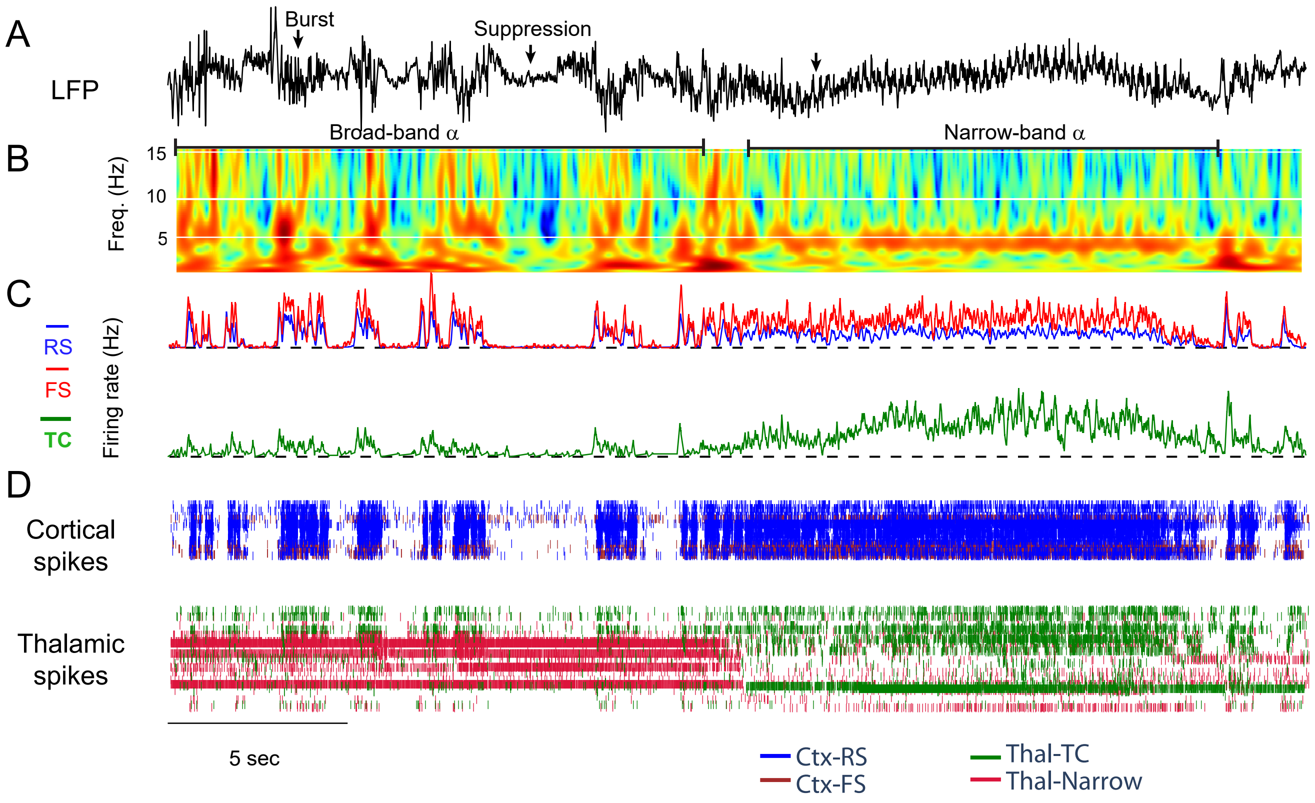


**Figure S6: LFP and raster plots during the transition from burst suppressions to narrow-band 5 Hz $\alpha$ oscillation.** **(A)** Raw LFP. **(B)** Scalogram. **(C)** Average firing rate of cortical and thalamic neurons. **(D)** Raster plot of cortical inhibitory ('Ctx-FS'), excitatory('Ctx-RS') neurons and thalamic excitatory ('Thal-TC') and thalamic inhibitory ('Thal-Narrow') neurons (see Methods).

### Figure S7



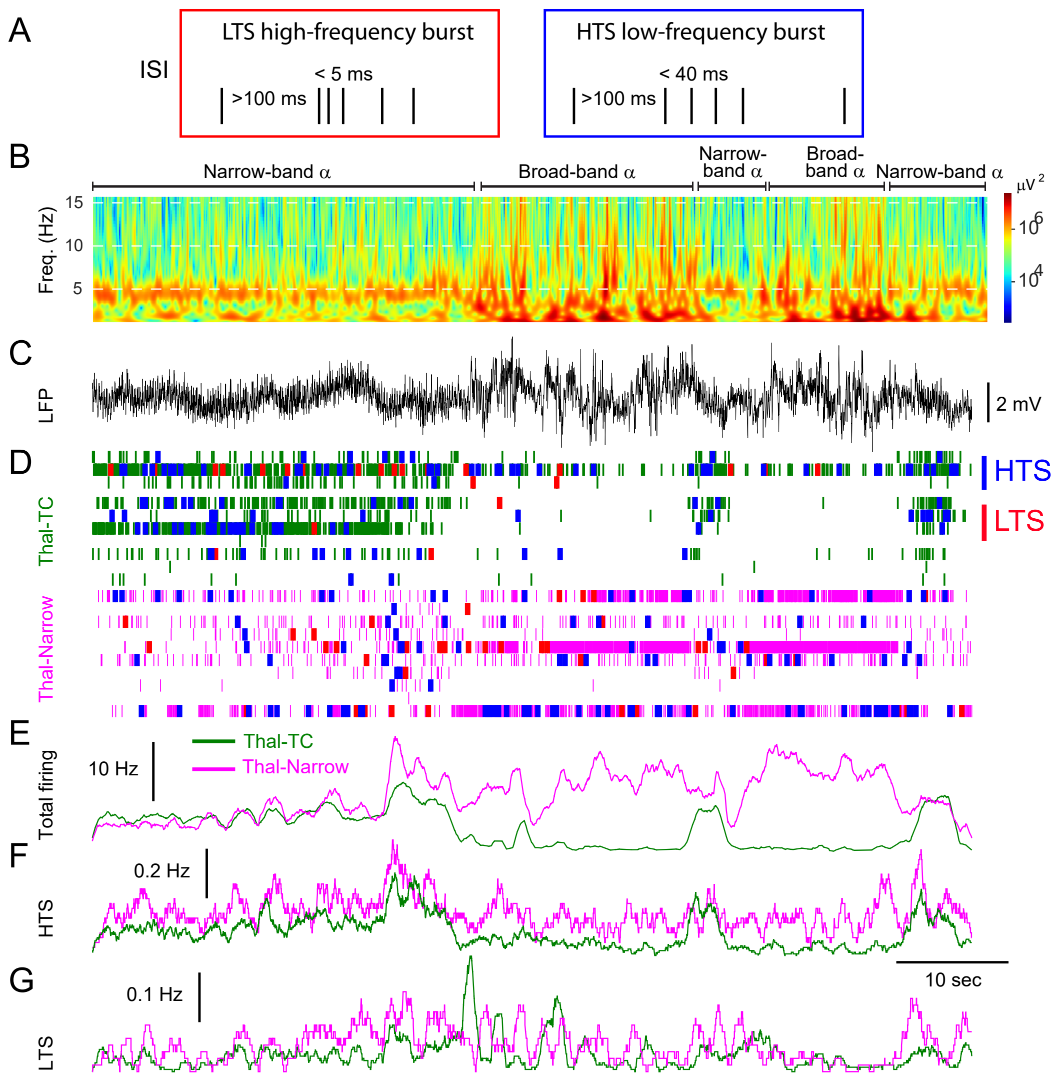


**Figure S7: Thalamic bursting at the transition from narrow-band 5 Hz to broad-band $\alpha$ oscillations.** **(A)** Definition of low-threshold spike (LTS) high-frequency bursts of action potentials: an LTS high-frequency burst is a group of action potentials a) preceded by a 100-msec interval with no action potentials and b) starts with an ISI between the first two action potentials $< 5$ msec. The definition of high-threshold spike (HTS) mediated burst applies to other groups of action potentials preceded by an interval of $> 100$ msec and starting by an interval of $< 40$ msec. **(B)** Scalogram of LFP. **(C)** Raw LFP. **(D)** Example raster plot. Top: 10 TC neurons showing the overlay of action potentials (green), LTS bursts of action potentials (red) and HTS bursts of action potentials (blue). Bottom same for narrow-spike thalamic neurons. Spikes are in magenta color. **(E)** Average firing rate of 161 TC neurons (green line) and of $n = 63$ inhibitory thalamic neurons (narrow spike)(magenta line) at the transition between narrow-band $\alpha$ oscillation and broad-band frequency $\alpha$ oscillations. **(F)** Same as in E but for HTS bursts **(G)** Same as in E but for LTS bursts. Note that the rate of TC neuron HTS bursts tends to decrease as narrow-band $\alpha$ oscillations disappear, whereas the rate of TC neuron LTS bursts remains relatively constant despite an overall decrease in TC firing rate.

### Figure S8



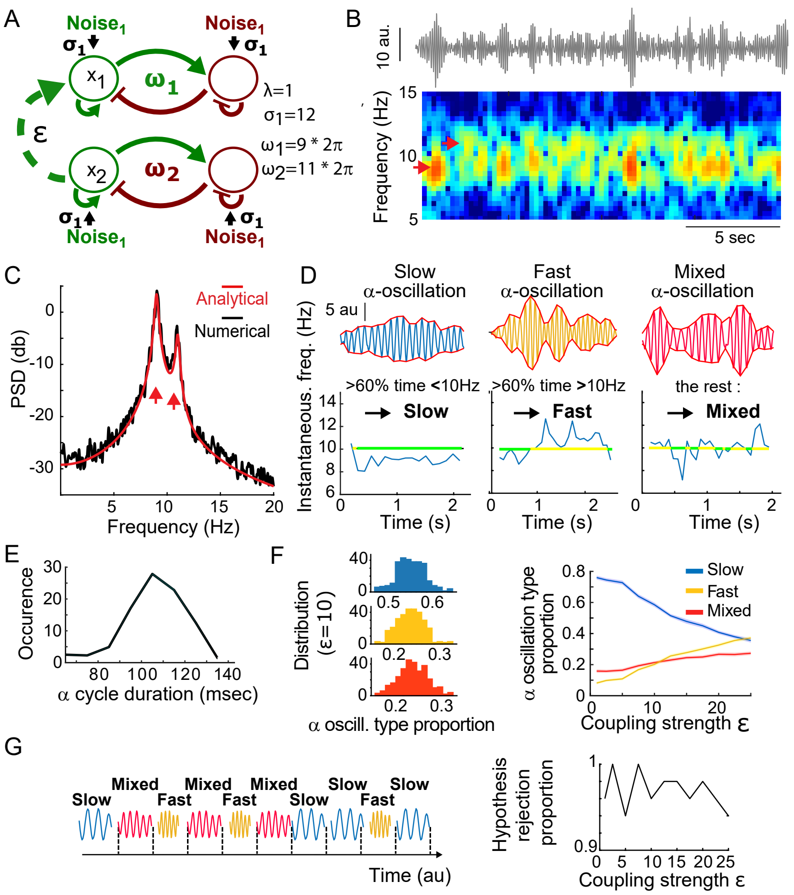


**Figure S8: $\alpha$ oscillations of mixed frequencies reflect two coupled 2-Dimension OU processes.** **(A)** Schematic model of two coupled 2D-OU processes and their parameters. **(B)** Realization with $\epsilon = 10$ (top) and the associated spectrogram (bottom). Local maxima in the color-coded spectrogram (red arrows) spread around 9 and 11 Hz. Amplitude and power are in arbitrary units ([a.u]. **(C)** Numerical (black) and theoretical (red) PSD (log scale) of the coupled 2D OU realization shown in (A) indicate 2 peaks (arrows). **(D)** Three segmented $\alpha$ oscillation (top) and their instantaneous frequency over time (bottom). The proportion of time spent with instantaneous frequencies below (green) or above (yellow) the mean frequency $(f_1 + f_2)/2 = 10Hz$ determines the classification into categories slow (more than 60% below 10Hz), fast (more than 60% above 10Hz) and mixed (the rest) $\alpha$ oscillations. **(E)** Distribution of the duration of the $\alpha$ oscillation cycle in panel B showing a continuum of cycle duration and thus a mixing of the two processes. **(F)** Left: distribution of $\alpha$ oscillations of categories for $\varepsilon = 10$ in 5-minute-long simulations indicates a stochastic process. Right: evolution of $\alpha$ oscillations of different frequencies with respect to the coupling strength $\varepsilon$ for fifty 5-minute-long simulations indicates an increase of fast $\alpha$ oscillations when the coupling strength increases. **(G)** Top: Schematic of time-sequenced $\alpha$ oscillations. Bottom: Statistical Runs test on fifty 5-minute-long simulations for each choice of $\varepsilon$, to determine the proportion of trials in which the null hypothesis (H0: the time sequence of oscillation types is deterministic) was rejected. The result indicate that the order is most of the time considered random as H0 is rejected (close to 1) for all tested $\varepsilon$. Thus $\alpha$ oscillation type is considered to appear in random order. See supplementary detailed results below.

#### Detailed results of Fig. S8

When $\alpha$ oscillations are in the broad-band regime with mixed $\alpha$ frequencies, we wondered whether they could be subject to some hierarchical properties of interactions. Therefore, we studied how the $\alpha$ frequency is distributed when two stochastic neural mass models are coupled together (eq. 1) (Fig. S8A) with a coupling parameter $\varepsilon = 10$. For that goal, we obtained the analytical expression (eq. 2) and computed the power spectral density of $\alpha$ oscillation generated by the model (Fig. S8B,C).
We further segmented the $\alpha$ oscillation and differentiated them into slow, fast, or mixed frequency based on their instantaneous frequency (Fig.S8D). The distribution of inverse instantaneous frequencies indicates a large distribution (Fig. S8E).
To further analyze the distribution of $\alpha$ oscillations, we then extracted the proportion of $\alpha$ oscillation: slow 0.55, fast 0.23 and mixed 0.22, which are broadly distributed (n=$33.10^3$ (Fig. S8F left). When increasing the coupling strength $\varepsilon$, the proportion of slow $\alpha$ oscillation decreases, while the mixed and fast ones increase (Fig. S8F right).
Finally, we calculate the probability of having two consecutive $\alpha$ oscillations of a given type. For that goal, we apply the conditional entropy $H_c(Y|X)$: and obtain $H_c = 1.45$ with 3772 $\alpha$ oscillations. We then uniformly reshuffled the $\alpha$ oscillation order 100 times from which the conditional entropy gave the same values, confirming that there was no deterministic order in the sequence generated by eq. 1.
Next, we studied whether the sequences of slow, fast, and mixed $\alpha$ oscillations were random or could appear with a specific deterministic order. To investigate whether $\alpha$ oscillation sequences carry out any information (Fig. S8G), we applied the Wald-Wolfowitz run test [1, 2] on 50 realizations of the coupled processes for 5 minutes to determine the proportion of trials where the null hypothesis H0 ("the sequence is deterministic") is accepted. The null hypothesis was rejected with a rejection rate $> 0.9$ (non-significant p-values for, respectively, 90 percent). This result supports the hypothesis that the sequence $\alpha$ oscillation of different frequencies — slow, fast and mixed - emerges in a statistically random order for a broad range of coupling strengths $\epsilon$. In addition, at least in 90% of the trials, the hypothesis of a deterministic order was rejected for all range of $\varepsilon$ in [0-25], confirming that there is no specific order in the appearance of the three types of $\alpha$ oscillations.

### Figure S9



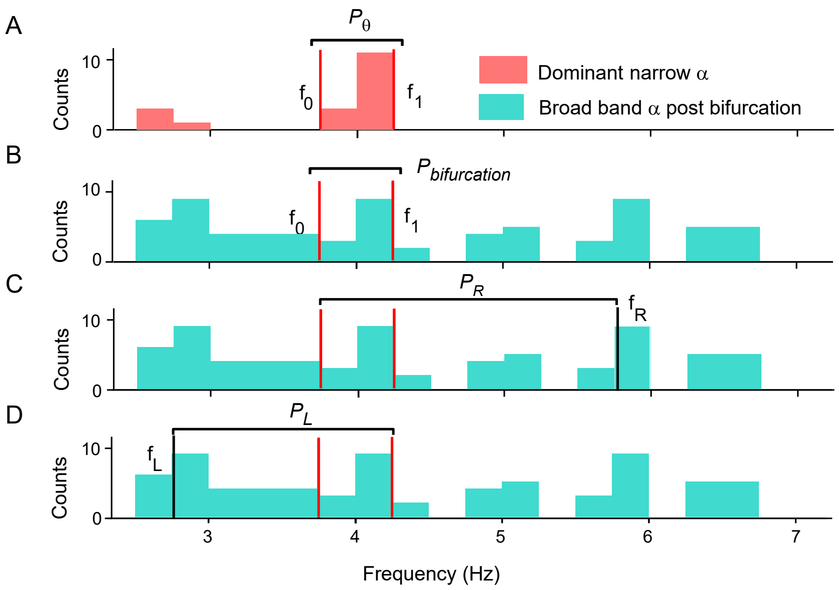


**Figure S9: Method of estimation of $\alpha$ oscillation frequency distribution before and after a bifurcation as in fig.7D. (A)** Distribution of individual narrow-band frequencies $[f_0, f_1]$ (defined as $\pm 0.25$ Hz around the peak) of $\alpha$ oscillations before the bifurcation time (Fig.7D). **(B)** After the bifurcation time, the spread of the frequencies outside the narrow band $[f_0, f_1]$ can go from 2.5Hz up to 6.75Hz in this example. **(C)** The estimation of the upper right frequency boundary $f_R$ corresponds to when the proportion between $f_1$ and $f_R$ of the selected $\alpha$ oscillations, noted $P_r$ reaches 0.75. **(D)** The estimation of the lower frequency boundary $f_L$ corresponds to when the proportion of oscillations between $f_L$ and $f_1$ of the selected $\alpha$ oscillation, noted $P_L$ reaches 0.75. See methods

### 2 Supplementary Materials and Methods

#### 2.1 Coupled Modeling

In a simpler coupling form, the equation of the system is as follows.

$$
\begin{aligned}
\dot{X}_1 &= -\lambda X_1 + \omega_1 Y_1 + \sqrt{2\sigma_1}\dot{w}_1 + \epsilon X_2 \\
\dot{Y}_1 &= -\omega_1 X_1 - \lambda Y_1 + \sqrt{2\sigma_1}\dot{w}_2 \\
\dot{X}_2 &= -\lambda X_2 + \omega_2 Y_2 + \sqrt{2\sigma_2}\dot{w}_3 \\
\dot{Y}_2 &= -\omega_2 X_2 - \lambda Y_2 + \sqrt{2\sigma_2}\dot{w}_4,
\end{aligned} \tag{1}
$$

#### 2.2 Power Spectral Density function for the coupled Eq.1

We present here the fit for the power spectrum density $S$ of Eq.1 using the exact formula:

$$
\begin{aligned}
S(f) = \frac{Z_\epsilon(\lambda,\omega_1,\omega_2)}{2\pi}&(A_\epsilon F_1(f;\omega_2) + B_\epsilon(\lambda,\omega_1,\omega_2)F_2(f;\omega_2) \\
&+ C_\epsilon(\lambda,\omega_1,\omega_2)F_3(f;\omega_1) + D_\epsilon(\lambda,\omega_1,\omega_2)F_4(f;\omega_1))
\end{aligned} \tag{2}
$$

where

$$
\begin{aligned}
F_1(f,\omega_2) &= \tfrac{\omega_2-2\pi f}{\lambda^2+(\omega_2-2\pi f)^2} + \tfrac{\omega_2+2\pi f}{\lambda^2+(\omega_2+2\pi f)^2} \\
F_2(f,\omega_2) &= \tfrac{\lambda}{\lambda^2+(\omega_2-2\pi f)^2} + \tfrac{\lambda}{\lambda^2+(\omega_2+2\pi f)^2} \\
F_3(f,\omega_1) &= \tfrac{\omega_1-2\pi f}{\lambda^2+(\omega_1-2\pi f)^2} + \tfrac{\omega_1+2\pi f}{\lambda^2+(\omega_1+2\pi f)^2} \\
F_4(f,\omega_1) &= \tfrac{\lambda}{\lambda^2+(\omega_1-2\pi f)^2} + \tfrac{\lambda}{\lambda^2+(\omega_1+2\pi f)^2}
\end{aligned}
$$

and

$$
\begin{aligned}
Z_\epsilon(\lambda,\omega_1,\omega_2) &= \frac{\sqrt{\sigma_1\sigma_2}}{2\lambda(\omega_2^2-\omega_1^2)\left(\lambda^2+\omega_1^2\right)\left((\omega_2-\omega_1)^2+4\lambda^2\right)\left((\omega_2+\omega_1)^2+4\lambda^2\right)} \\
A_\epsilon(\lambda,\omega_1,\omega_2) &= 4\omega_2\lambda\epsilon^2\left(\lambda^2+\omega_1^2\right)\left(\omega_2^2+4\lambda^2+\omega_1^2\right) \\
B_\epsilon(\lambda,\omega_1,\omega_2) &= 2\omega_2^2\epsilon^2\left(\lambda^2+\omega_1^2\right)\left(\omega_2^2+4\lambda^2-\omega_1^2\right) \\
C_\epsilon(\lambda,\omega_1,\omega_2) &= -\lambda\omega_1\epsilon^2\left(\omega_2^4+6\omega_2^2\omega_1^2+16\lambda^4+8\lambda^2\left(2\omega_2^2+\omega_1^2\right)+\omega_1^4\right)
\end{aligned}
$$

$$
\begin{aligned}
D_\epsilon(\lambda,\omega_1,\omega_2) &= -\omega_1^6(-6\omega_2^2+18\lambda^2+\epsilon^2) \\
&+ 2\omega_2^2\lambda^2(\omega_2^2+4\lambda^2)(\omega_2^2+4\lambda^2+\epsilon^2) \\
&\quad -2\omega_1^4(3\omega_2^4+24\lambda^4+\lambda^2(4\epsilon^2-3\omega_2^2)) \\
&+ \omega_1^2(\omega_2^4(2\omega_2^2+\epsilon^2)-32\lambda^6+16\lambda^4(2\omega_2^2-\epsilon^2) \\
&+ 2\lambda^2(5\omega_2^4-\omega_2^2\epsilon^2))-2\omega_1^8.
\end{aligned}
$$

### References

[1] J. V. Bradley, *Distribution-free statistical tests*, vol. 60. United States Air Force, 1960.

[2] D. J. Sheskin, *Handbook of parametric and nonparametric statistical procedures*. Chapman and hall/CRC, 2003.
